# Deep Recovery of Effective Migration Surfaces

**DOI:** 10.64898/2026.11.03.763937

**Authors:** Jiatong Liang, Gideon Bradburd, Jonathan Terhorst

## Abstract

The isolation-by-distance (IBD) principle describes the tendency for genetic relatedness to decline with geographic distance. Although the existence IBD has long been documented in natural populations, recent analyses of spatially referenced genotype data have shown that the strength of this relationship can vary across a landscape due to geographic barriers, habitat compatibility, or other factors. Effective migration surfaces (EMS) have emerged as a popular exploratory tool for visualizing spatial patterns in IBD. However, fitting likelihood-based EMS models is often computationally demanding, requiring repeated large-scale matrix calculations. Here, we introduce Deep Recovery of Effective Migration Surfaces (dreems), a graph neural network for amortized inference of effective migration surfaces. dreems maps genetic covariance and sampling locations directly to an effective migration surface, while simultaneously providing predictive uncertainty estimates. Once trained, the network accommodates different graph sizes and sampling designs without needing to re-train. Across independent coalescent simulations that were not seen during training, dreems has roughly 10-30% lower estimation error compared to a state-of-the-art likelihood-based method, with the largest gains observed under sparse sampling designs that are commonly encountered in practice. Reanalysis of several published datasets recovers spatial patterns that are broadly consistent with earlier likelihood-based estimates, while reducing computation time by orders of magnitude.

## 1 Introduction

Understanding where and how organisms disperse has important applications across many areas of ecology and evolutionary biology. For example, in conservation, connectivity between populations can guide the placement of protected areas (Gaines et al., 2010; Wilcox et al., 2023). In epidemiology, identifying barriers and routes for the movement of disease vectors can help anticipate and limit their spread (Dong et al., 2025; Tatem et al., 2006). Dispersal and gene flow also shape demographic history (Bradburd and Ralph, 2019; Slatkin, 1987) and the geographic distribution of adaptive variation (Lenormand, 2002).

Classical models of spatial population structure predict that when dispersal is local and spatially uniform, genetic differentiation tends to increase with geographic distance (Malécot, 1948; Wright, 1943), a relationship which Wright termed *isolation-by-distance* (IBD). Subsequent theoretical work showed that geographic barriers and spatial variation in dispersal and population density can alter the relationship between genetic differentiation and geographic distance (Ayati et al., 1999; Nagylaki, 1988; Nagylaki et al., 1993).

Increasing access to data and computation now enable us to directly observe these patterns in nature (e.g., Jay et al., 2013; Twyford et al., 2020; Wilcox et al., 2023). An especially popular visualization technique is the effective migration surface (EMS) (Marcus et al., 2021; Petkova et al., 2016), which represents a study region as a graph, with nodes corresponding to demes and edges to potential migration pathways. Edge weights indicate greater or less connectivity between adjacent demes, relative to an overall background rate of spatial connectivity (or migration). When overlaid atop a map, the EMS highlights regions of relatively high or low connectivity, helping to identify potential barriers to gene flow, migration corridors, or other features.

Most existing methods for estimating EMS are likelihood-based: they specify a probability model for genetic data or summary statistics and fit its spatial parameters to the dataset. Estimating Effective Migration Surfaces (EEMS) uses Bayesian inference with an electrical-resistance approximation to expected genetic dissimilarity (Petkova et al., 2016). In this approximation, metapopulations are modeled as electrical circuits for which the “resistance” between two nodes accounts for connectivity through all paths that connect them (McRae, 2006). There have been a number of extensions of the original EEMS approach, including Fast Estimation of Effective Migration Surfaces (feems), which provides efficient penalized-likelihood estimation using a Gaussian Markov random field formulation (Marcus et al., 2021), and Migration and Population-size Surfaces (MAPS), which uses a coalescent model of identity-by-descent sharing to estimate migration and population-size variation over recent time periods (Al-Asadi et al., 2019). feems and MAPS have been extended in turn, generating feemsmix, which adds directed long-range ancestry connections in addition to local migration (Shastry et al., 2025), and Fine-Resolution Asymmetric Migration Estimation (FRAME), which allows for asymmetric migration between neighboring populations (Shen and Novembre, 2026).

Fitting these models can be computationally demanding because likelihood evaluation requires recalculating the genetic relationships implied by each candidate set of migration and population-size parameters. In graph-based models, these calculations involve matrix operations whose cost grows at least cubically in size of the spatial grid (Shen and Novembre, 2026). Model fitting, posterior sampling, or hyperparameter tuning may invoke the likelihood function thousands of times and require hours of computing time.

Difficulty evaluating likelihoods has a long history in population genetic inference, to the point where a whole literature on “likelihood-free” inference methods has emerged. In place of a likelihood function, these methods rely on simulations to assess how well parameters fit a given dataset. An early example is approximate Bayesian computation (ABC), which approximates a posterior distribution by comparing simulated and observed summary statistics (Beaumont et al., 2002). More recently, machine learning and neural methods trained on simulated data have greatly extended the scope of likelihood-free inference in population genetics (Huang et al., 2024; Schrider and Kern, 2018). Noteworthy applications include detection of selective sweeps (Riley et al., 2024; Schrider and Kern, 2016; Sugden et al., 2018; Whitehouse and Schrider, 2023), demographic inference (Flagel et al., 2019; Sheehan and Song, 2016; Wang et al., 2021), recombination rate estimation and hotspot detection (Adrion et al., 2020; Chan et al., 2018), and estimation of coalescence times and variant ages (Nait Saada et al., 2023). A particular appeal of many of these methods is that they are *amortized* : the large initial cost of simulation and training is spread across subsequent analyses, and the trained network can be applied to new datasets without retraining.

Here, we introduce Deep Recovery of Effective Migration Surfaces (dreems), a graph neural network (GNN) that maps genetic covariance to relative migration rates, local coalescence rates, and predictive uncertainty. Its architecture combines local and graph-wide information and accommodates different graph sizes and sampling designs without the need to re-train. Compared with a state-of-the-art likelihood-based method (Marcus et al., 2021), dreems achieves higher mean accuracy on unseen datasets, particularly ones with limited samples. Furthermore, dreems produces estimates in seconds. Reanalysis of several published datasets shows that dreems largely reproduces earlier, likelihood-based estimates, with some potentially new findings arising from its specialization to sparse data and ability to explore more complex graph structures.

The rest of this paper is organized as follows. Section 2 describes the dreems archi-tecture, covariance refinement, training, and uncertainty calibration. Section 3 presents simulation benchmarks and empirical applications, including sensitivity to distribution shift, grid design, and ploidy. Section 4 discusses limitations and possible extensions.

## 2 Methods

### 2.1 Overview of the dreems architecture

Figure 1 summarizes how dreems maps genetic covariance, sample coordinates, and a user-specified spatial graph *G* = (*V, E*) onto an effective migration surface. The graph has *n* = *V* nodes representing demes and *m* = |*E*| undirected edges representing potential migration between adjacent demes. Each edge has a positive weight *w*_*e*_, with larger values representing greater effective migration. Samples are assigned to their nearest nodes. Let *N*_*i*_ be the number of individuals assigned to node *i*, |*O*| = {*i* : *N*_*i*_ > 0} the set of observed For *N* = Σ_*i*_*N*_*i*_ individuals and *s* SNPs, let *X* ∈ R^*N* ×*s*^ contain genotype dosages. We demes, and *o*= |*O*| their number; the remaining *n* − *o* demes are unobserved.

**Figure 1:**
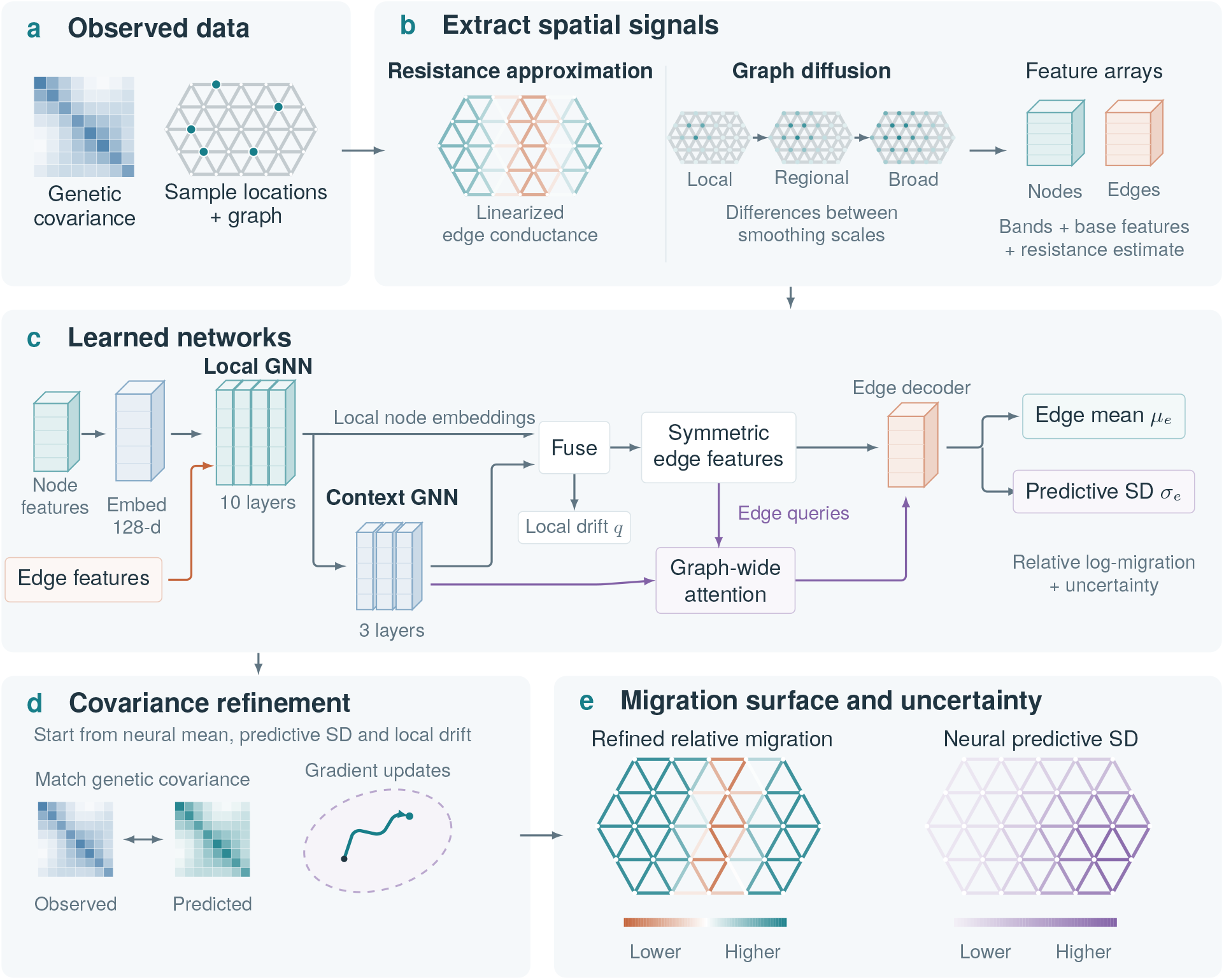
Overview of dreems. **(a)** Sample locations assign genetic covariance to demes on a user-supplied graph. **(b)** A linearized resistance estimate and six diffusion bands of covariance and sampling signals provide edge and node features. **(c)** A network trained on simulated spatial histories combines message-passing layers and graph-wide edge-to-node attention. It predicts a mean relative log-migration rate and predictive standard deviation (SD) for each edge, together with node-level local coalescence rates. **(d)** Additional optimization steps improve the covariance fit. **(e)** The refined migration surface is reported alongside the neural predictive SD.

For *N* = Σ_*i*_ *N_i_* individuals and *s* SNPs, let *X* ∈ R^*N*×*s*^ contain genotype dosages. We center each SNP across individuals and divide by its sample standard deviation to obtain *Y*. The individual-level genetic covariance is *C*^ind^ = *Y Y* ^T^*/s*. Let *T* ∈ R^*o*×*N*^, with rows indexed by *O*, average individuals within each observed deme: *T*_*ia*_ = 1*/N*_*i*_ if individual *a* is assigned to deme *i*, and zero otherwise. Averaged within demes, we define the observed covariance

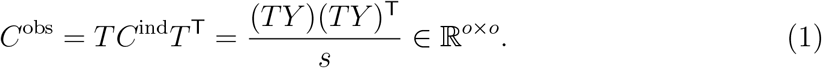

Thus, each entry averages genetic covariance over all individual pairs in the corresponding demes, including self-pairs for diagonal entries. To featurize *C*^obs^, we embed it in an *n* × *n* matrix *C*, placing zeros in rows and columns for unobserved demes. These zeros encode missing observations, rather than measured absence of genetic covariance, and the neural network is also given sample counts and observation indicators to identify which demes were sampled.

Before applying the neural network, dreems summarizes the genetic data and sampling design as features attached to graph nodes and edges (Figure 1b). Node features describe genetic variation at each deme and the amount of data supporting it. Edge features describe contrasts between neighboring demes and evidence about their connectivity. We construct these features in two ways: first, by estimating a rough migration surface from resistance distances, and second, by summarizing genetic variation at several spatial scales. These correspond to the “All-pairs resistance” and “Graph diffusion” calculations in Figure 1b, respectively.

The first set of features provides a cheap initial migration estimate via a linear approximation to resistance distances. We suppose observed genetic dissimilarity is approximately proportional to the effective resistance distance between the corresponding demes: *D*_*ij*_ ≈*cR*_*ij*_(***θ***), where *D*_*ij*_ = *C*_*ii*_ + *C*_*jj*_ −2*C*_*ij*_ is calculated from the deme-wise covariance matrix *C, c >* 0 is a common scale factor, and *θ*_*e*_ = log *w*_*e*_ is the log-weight of edge *e*. Taking logarithms gives log *D*_*ij*_ ≈ log *c* + log *R*_*ij*_(***θ***). On a connected graph with positive edge weights, log *R*_*ij*_ is a smooth function of ***θ***. We write ***θ*** = ***θ***^(0)^ + ***δ***, where *θ*_*e*_^(0)^ = log *w*_*e*_^(0)^. The baseline edge weights are initialized from edge lengths, with shorter edges receiving greater weight. By Taylor expansion as ***δ***→ **0**,

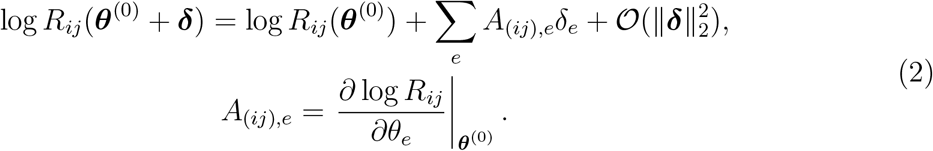

We then have

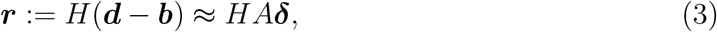

where *H* = *I*_*p*_ −**11**^T^*/p* is a centering matrix that removes the unknown common scale log *c*. The normal equations for (3) involve the matrix

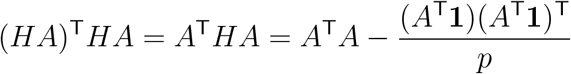

which is potentially high-dimensional. For a fast initialization, we neglect the rank-one term and fit only the diagonal of *A*^T^*A*, and add ridge regularization and smoothing between neighboring edges to stabilize the estimate. With *a*_*e*_ =Σ_*i*<*j*_ *A*^2^ _(*ij*),*e*_, we solve

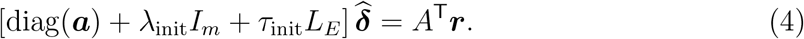

Here *L*_*E*_ is the unweighted Laplacian of a line graph whose nodes represent migration edges and edges connect when those edges share a deme. We use *τ*_init_ = 0.01 and set *λ*_init_ to 10^−3^ times the median positive *a*_*e*_, with a numerical floor of 10^−12^. The resulting edge field is centered and scaled to unit SD for input to the network.

We also feed transformed principal-component scores from *C*^obs^ into the network. Let (*λ*_*k*_, ***v***_*k*_) denote its eigenpairs, ordered from largest to smallest eigenvalue. An observed node *i* receives 20 coordinates,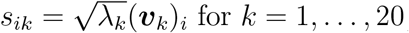 for *k* = 1, …, 20, as part of its input to the network. We choose each eigenvector’s sign so that its largest absolute entry is positive. Scores are zero at unobserved nodes, and unused coordinates are zero when fewer than 20 components are available.

For graph diffusion, we form a matrix *S* ∈ ℝ^*n×*13^ with one row per node. Its first eight columns reuse the first eight of these 20 PCA coordinates. The remaining five columns contain:

1. The sample count, entered as log(1 + *N*_*i*_).
2. An observation indicator, equal to one if *N*_*i*_ > 0 and zero otherwise.
3. Summaries of the precision of each entry of the covariance matrix:
4. For the diagonal entries *C*_*ii*_, it equals the number of individual pairs within deme *i* divided by an estimate of the sampling variance of *C*_*ii*_.
5. For off-diagonal entries *C*_*ij*_ with *i* ≠ *j*, it equals the number of individual pairs between demes *i* and *j* divided by an estimate of the sampling variance of *C*_*ij*_.

These summaries are set to zero at unobserved nodes. Additionally, we log-transform the precisions using *u* →log(1 + *u*), and rescale the nonzero values to have maximum one for input into the network.

Finally we standardize each of the 13 columns across graph nodes and clip them to [−8, 8], and then diffuse these signals over the graph (Figure 1b). Diffusion uses the normalized graph Laplacian 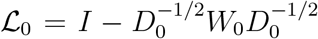, where *W*_0_ is the symmetric adjacency matrix containing the baseline weights 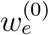, and *D*_0_ = diag(*W*_0_**1**) is the degree matrix. We then compute

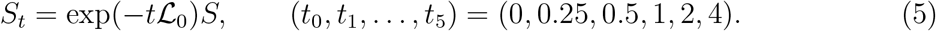

The matrix exponential suppresses graph modes with larger Laplacian eigenvalues more strongly as *t* increases. As input to the neural network, we pass changes between consecutive smoothing scales:

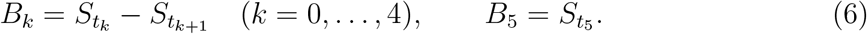

These six matrices sum to *S*, separating variation removed at successive scales from the remaining broad-scale component (Figure 1b).

The preceding steps result in 6 × 13 = 78 diffusion features ***z***_*i*_ for each node, which we append to its 20 PCA coordinates and sampling features. To compute edge features, we assign to edge (*i, j*) three symmetric summaries of its endpoints’ features, with all operations applied elementwise:

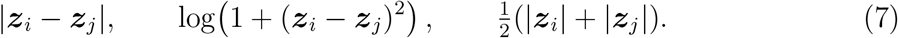

These provide 3 × 6 ×13 = 234 edge features describing endpoint contrasts and signal magnitudes. Together with the resistance estimate and the base genetic and sampling features, these form the node and edge “Feature arrays” at the right of Figure 1b, which enter the neural network in panel c.

A learned input layer (“Embed 128-d” in Figure 1c) maps each node’s feature vector to an initial 128-dimensional state for message passing. A local graph neural network (GNN) block then updates each node using information from its neighbors. In each layer, the edge features use a learned gate that weights the neighboring node’s channels, together with an additive message describing the connecting edge. The receiving node averages these messages and combines them with its current representation. Each update is added to the previous representation, forming a residual connection that retains earlier information. This operation is repeated ten times, allowing each node to incorporate evidence from progressively more distant neighborhoods. Additionally, we pass the resulting messages through three additional layers that average the node states without the edge-dependent gates used in the first ten layers in order to aggregate messages at different depths, which has previously been shown to improve learning (Xu et al., 2018). Finally, we fuse the states before and after these three layers and apply a learned transformation to produce the final node embeddings.

For each edge, the network then takes the sum and absolute difference of its two endpoint messages, and appends the original edge features. This forms the “Symmetric edge features” in panel c; swapping the endpoints leaves these features unchanged, consistent with our inductive bias that migration is symmetric between nodes.

The “Graph-wide attention” block in Figure 1c enables each edge to use information beyond the neighborhood reachable by local message passing. Each edge message is used to produce a learned query, and every node message produces a key and a value. An attention mechanism then assigns a normalized weight to each node, and the weighted average of the values provides a graph-wide summary specific to that edge. These are represented by the “Edge queries” and “All nodes: keys / values” paths in panel c. Thus, different edges can draw on different distant regions. Finally, the “Edge decoder” combines this summary with the edge’s endpoint and input features to produce the “Edge mean” and “Predictive SD” outputs.

Separate output layers predict the relative log-migration rate, *µ*_*e*_, and its predictive standard deviation, *σ*_*e*_, for every edge. Edge predictions are mean-centered, so exp(*µ*_*e*_) expresses migration relative to the geometric mean across edges. The final output is also passed through two additional learned gates: an edge-specific gate blends the decoder’s prediction with the resistance initializer, and a graph-level gate shrinks the centered field toward homogeneous migration when the graph summaries provide little evidence of spatial structure. See Supplementary Section S1.3 for further details. A separate layer predicts centered log-coalescence rates log *q*_*i*_ at nodes. Higher *q*_*i*_ corresponds to smaller local effective population size. The edge SD *σ*_*e*_ describes predictive uncertainty in centered log-migration. Its calibration is evaluated in Section 3.

### 2.2 Edge refinement

The neural network is primarily trained to minimize squared loss for relative migration rates. However, this does not necessarily imply that the predicted genetic covariance will match the observed covariance, particularly for ill-posed problems (e.g., when sampling is sparse). To improve concordance with the primary observable, we perform an edge refinement step by default (Figure 1d), which adjusts the migration weights to better match the observed genetic covariance. When developing our method, we observed that this refinement step tended to decrease error and increase mean correlation with the truth, so it is enabled by default and was used for all the fits reported below.

We approximate genetic covariance by a resistance-based spatial component *K*(***θ***) and a local residual component *Q* (Marcus et al., 2021). Let *P* = *I*_*o*_ − **11**^T^*/o* center the observed demes. The centered empirical covariance is 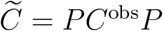. Our refinement procedure tries to minimize the distance between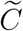 and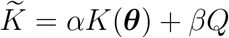, where

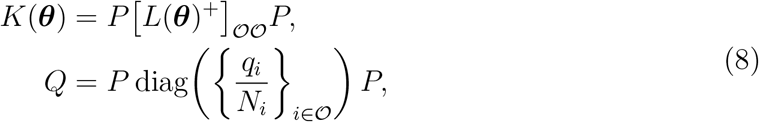

Here *L*(***θ***) is the graph Laplacian with edge weights exp(*θ*_*e*_), and *L*^+^ is its pseudoinverse. For *Q*, we use the predicted coalescence rates *q*_*i*_ as proxies for local variance, and divide by *N*_*i*_ to account for averaging individuals within a deme. The nonnegative coefficients *α* and *β* are fitted parameters that determine the scales of the two components.

We minimize covariance mismatch plus a small smoothness penalty, subject to a trust-region bound around the neural estimate:

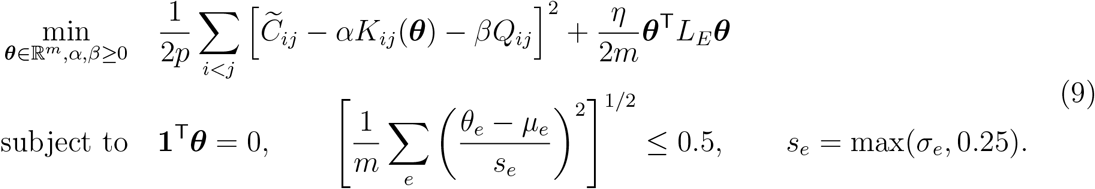

Here *η* = 10^*−*4^ imposes a small amount of smoothing between edges that share a deme. Starting from ***µ***, we take 50 gradient steps, refitting *α* and *β* at each step. The coalescence predictions and neural SDs remain fixed. Constraining the refined estimates in this way allows the network to scale the corrections relative to the amount of uncertainty in the neural predictions. This correction prioritizes sparse data problems, which are mostly encountered in practice. However, it can limit recovery when dense data support larger changes (Section 3.3).

### 2.3 Training on simulated data

Training data were generated using a custom structured coalescent simulator that is optimized for the stepping-stone model (Supplementary Section S1.1.1). The output of each simulation run is single biallelic SNP, generated under an assumed infinite sites model. Because private and low frequency mutations contribute little migration signal, we follow earlier work (Marcus et al., 2021; Petkova et al., 2016) and condition each SNP to have minor allele frequency (MAF) of at least 0.05 via rejection sampling. Each SNP is generated independently, which is equivalent to assuming that all SNPs are in linkage equilibrium.

To make the training data more realistic, we introduce variability in both the spatial structure and the demographic parameters across simulations. Training graphs are drawn from five “motifs:” real-world landmasses discretized on hexagonal lattices; synthetic corridors and coastlines; synthetic landmasses of varying sizes and shapes; randomly masked grids containing islands, bays, or lakes; and a simple lattice with fixed topology. These motifs are illustrated in Supplementary Figure S1. Further details about their construction and distribution are described in Supplementary Section S1.1.

Demographic parameters and sampling designs also vary randomly across simulations. Spatially correlated log-migration and log-coalescent rate fields are generated from Gaus-sian processes, with additional scenarios imposing homogeneous rates, discrete barriers, or habitat effects. Sampling designs include uniform selection, spatial transects, and clusters, with variable numbers of diploids at each sampled deme. Further details are given in Supplementary Section S1.2.

The network is trained to predict the centered log-rates. Its edge loss is Gaussian negative log-likelihood, which jointly trains the mean and SD:

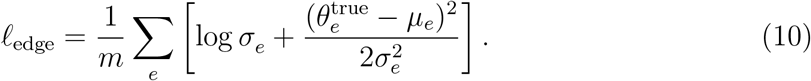

Additional loss terms are also added to improve training stability and generalization. For additional details about training see Supplementary Section S1.4.

### 2.4 Calibration of predictive uncertainty

To calibrate the predictive uncertainty of the network, we used a separate set of simulated datasets. After training, we split an additional 512 independently simulated datasets into approximately equally sized train, test, and validation sets. On the fitting set, we divided each edge’s error in the predicted centered log-migration rate by its raw predicted SD and averaged the squared ratios within each graph. The square root of the average across graphs gave a multiplicative SD correction, with each graph weighted equally. We also fitted separate corrections for graphs with at most 10%, more than 10% but at most 25%, or more than 25% of their nodes sampled. Before taking square roots, the average for each sampling group was combined with the overall average, giving the latter a weight equivalent to 25 graphs to stabilize the estimates. The selection set compared unchanged SDs, a single global correction, and corrections by sampling density, using the lowest Gaussian negative log-likelihood averaged over graphs. Network weights and predicted means remained fixed throughout. On the verification set, we checked how often Gaussian prediction intervals at nominal levels of 50%, 80%, and 95% contained the true centered log-migration rates.

## 3 Results

### 3.1 Performance on coalescent simulations

We compared dreems with feems (Marcus et al., 2021) on 512 newly generated coalescent datasets that were not seen during training. Accuracy was measured by Pearson correlation and mean squared error (MSE) between inferred and true edge log-migration rates, after centering each surface to remove its overall scale. We averaged the per-dataset correlations and MSEs, giving each dataset equal weight. The graphs in this validation set ranged from 219 to 1,406 nodes with 590 to 4,065 edges and a median of 39 sampled nodes. Each simulated dataset has a randomized migration and coalescent surface, sampling scheme, and graph design. A MAF filter of 0.05 was applied across all datasets. feems was evaluated on the identical set of inputs, with hyperparameter tuning performed using leave-on-out cross validation (LOOCV) over ten *λ* ∈ [10^*−*3^, 10^2^] and five *λ*_*q*_ ∈[10^*−*2^, 10^2^], as recommended by the package documentation.

Note that although these examples were not part of the training corpus, they are sampled from the same distribution of graphs and migration fields used to generate the training data. Section 3.3 evaluates model performance under a stronger form of distribution shift.

Across the validation set, mean Pearson correlation was 0.717 for dreems and 0.614 for feems; mean MSE was 0.539 and 0.738, respectively. The mean paired differences were 0.103 higher correlation and 0.199 lower MSE for dreems (Figure 3). The differences were largest under sparse sampling. Stratifying by sample design (Figure 2), dreems had a mean correlation 0.112 higher and MSE 0.211 lower when at most 10% of nodes were sampled (430 datasets). These differences narrowed to 0.061 and 0.143 for sampling of more than 10% but at most 25% of nodes (73 datasets), and 0.031 and 0.089 for sampling of more than 25% (nine datasets).

**Figure 2:**
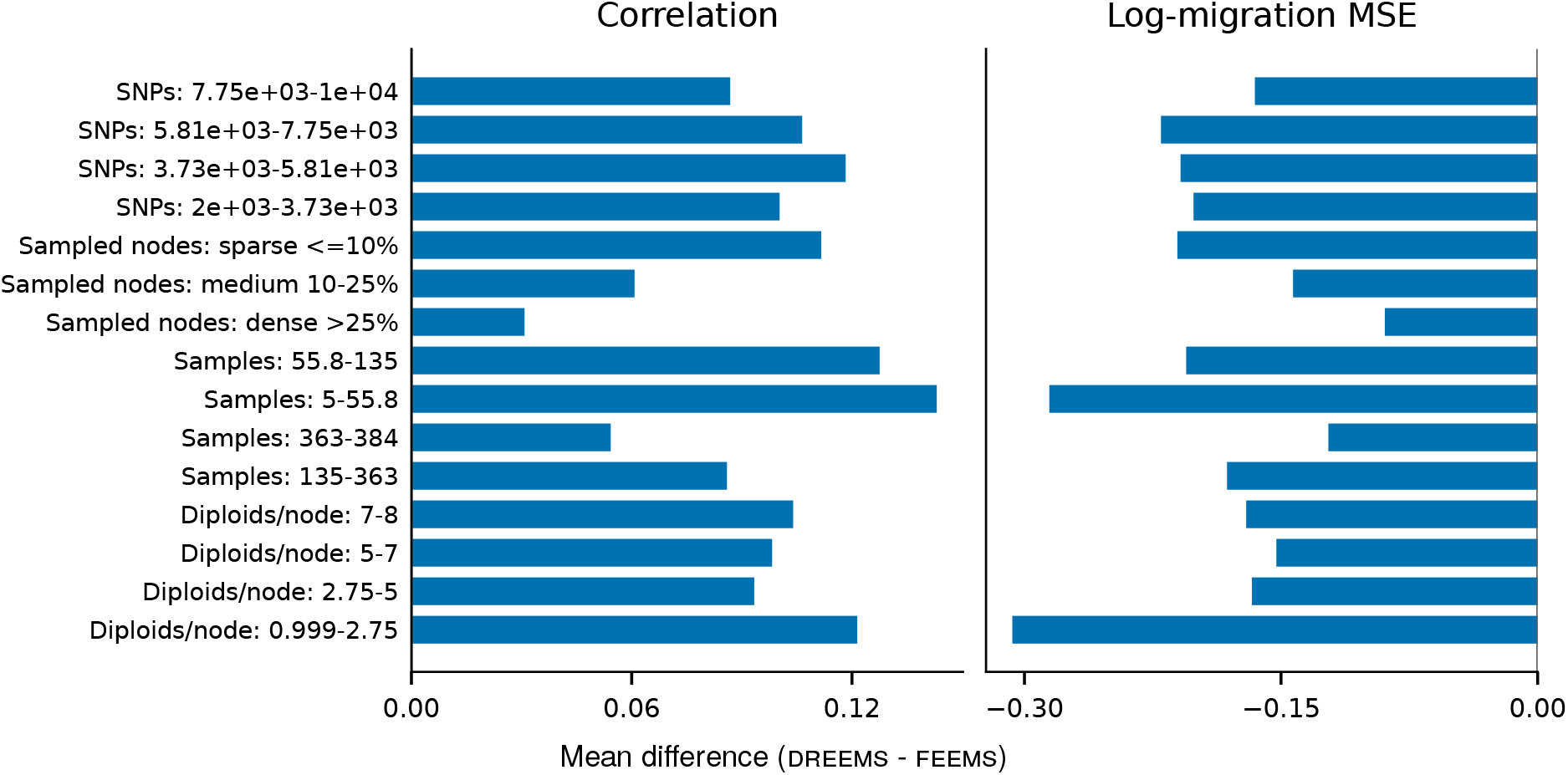
Accuracy differences between dreems and feems across SNP counts, sampling densities, total sample counts, and diploids per sampled node in the 512-dataset benchmark. Bars show mean dreems-minus-feems differences in Pearson correlation (left) and MSE of centered log-migration rates (right). Positive correlation differences and negative MSE differences favor dreems.

**Figure 3:**
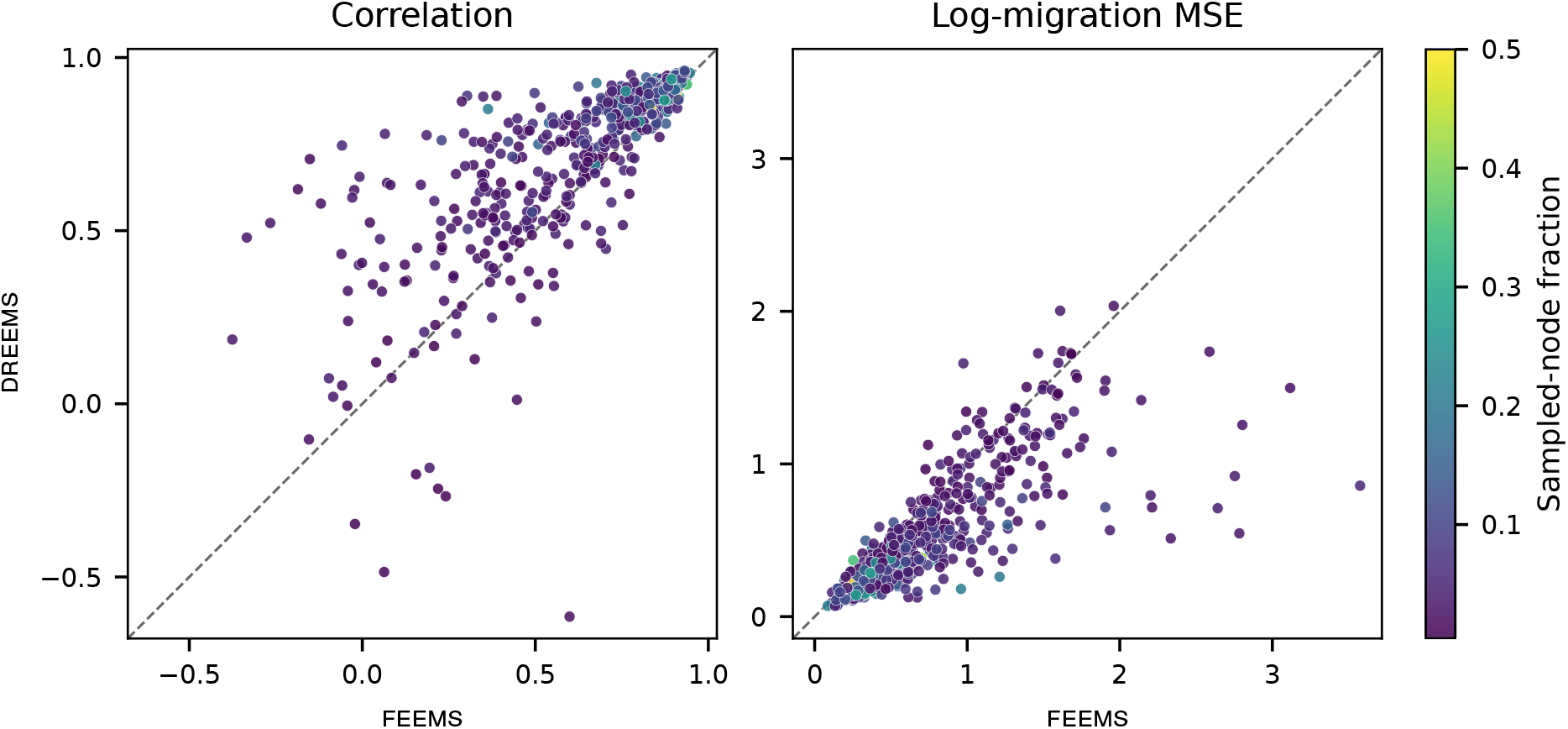
Comparison of dreems with feems tuned by leave-one-out cross-validation (LOOCV) across 512 randomly simulated migration landscapes. Each point represents one dataset; color indicates the fraction of nodes sampled. Points above the identity line in the correlation panel (left) or below it in the MSE panel (right) favor dreems.

dreems also had higher mean accuracy across the SNP-count and sample-size groups shown in Figure 2. In the lowest SNP-count quartile (2,003–3,727 SNPs), mean correlation was 0.100 higher and MSE 0.201 lower. In the lowest total-sample-size quartile (5–55 individuals), the corresponding differences were 0.143 and 0.285; in datasets averaging one to 2.75 diploids per sampled node, they were 0.121 and 0.307. The densely sampled datasets in this benchmark also favored dreems on average, although this group was small relative to the other two.

### 3.2 Predictive uncertainty

dreems predicts an SD for each edge’s centered log-migration rate (Section 2.1), with post-training calibration described in Section 2.4. On an independent verification split, mean coverage was 53.1%, 82.0%, and 94.8% at nominal levels of 50%, 80%, and 95% (Figure 4).

**Figure 4:**
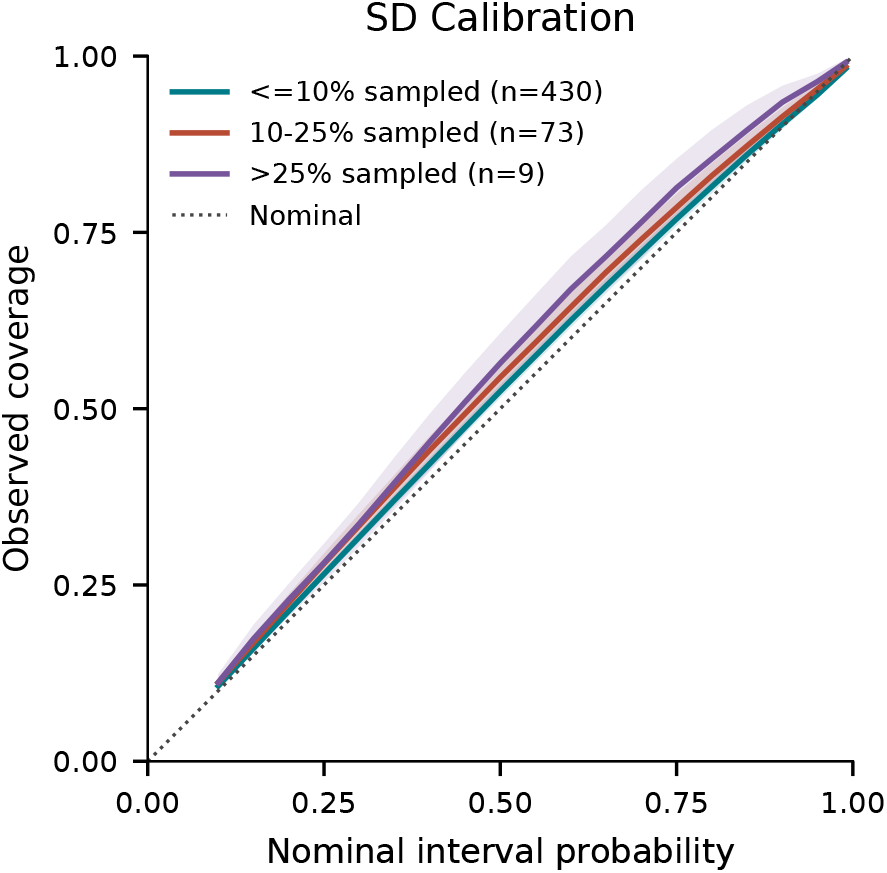
Marginal coverage of Gaussian prediction intervals around neural centered log-migration estimates before refinement, grouped by the fraction of graph nodes sampled. Graphs receive equal weight; shading shows pointwise 95% bootstrap intervals from resampling graphs. The dotted line indicates nominal coverage.

**Figure 5:**
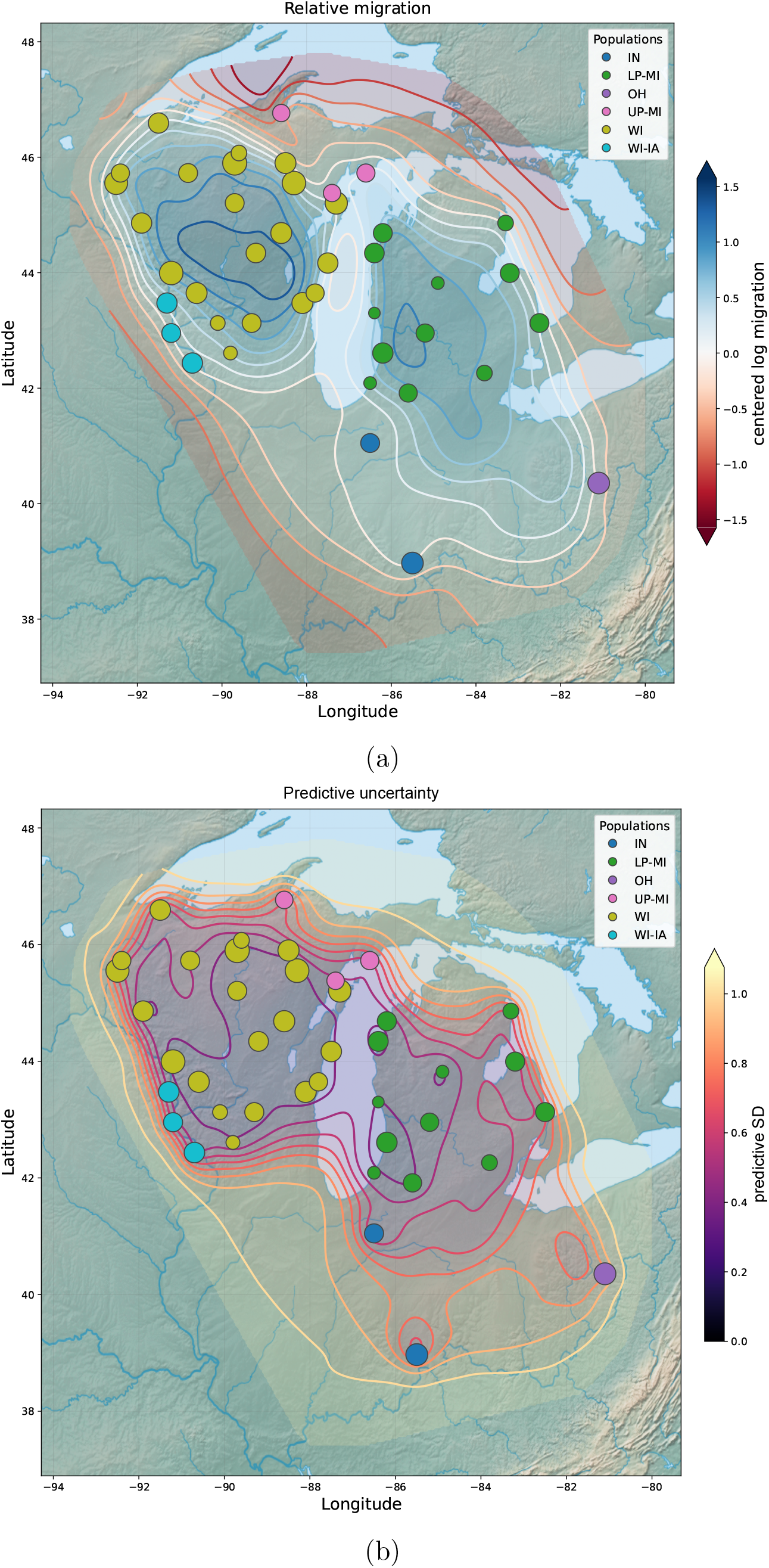
dreems estimates for blacklegged ticks on a grid with 2,860 nodes. **(a)** Relative log-migration, with blue indicating higher migration and red lower migration. **(b)** Neural predictive SD, with larger values indicating greater uncertainty. Node sizes indicate sample counts.

### 3.3 Assessing out-of-distribution performance

To assess performance under a strong distribution shift, we conducted a pilot study using migration surfaces generated by a Matérn-1*/*2 field. This field uses the kernel *k*(*d*) = exp(−*d/l*), where *d* is spatial distance and *l* is the correlation length scale. Compared to the squared-exponential kernel used in training, *k*(*d*) = exp[−*d*^2^*/*(2*l*^2^)], Matérn-1*/*2 surfaces are rougher and have nondifferentiable sample paths and greater variation at short distances. On a 10 ×10 triangular lattice, we drew four independent surfaces at each of three length scales (*l* = 0.12, 0.35, and 0.65), with node coordinates centered and scaled to unit maximum radius. Log-migration rates were obtained by averaging node fields over edge endpoints and standardizing to mean zero and unit variance. Each of the 12 surfaces was evaluated with nested sampling of 10% and 50% of nodes, four diploids per sampled node, and 2,000 SNPs, yielding 24 datasets. We applied the pretrained dreems model without retraining, using the same refinement settings and feems LOOCV protocol as above.

Across all these out-of-distribution test datasets, dreems’s gains in accuracy were not as substantial as those described above; dreems achieved an average correlation of 0.789, and mean MSE of 0.442, compared to a mean correlation of 0.744 and mean MSE of 0.550 for feems (Supplementary Figure S3). Additionally, we found that performance depended strongly on sampling density: dreems had lower MSE than feems in 11 of 12 sparse datasets (mean MSE 0.548 versus 0.888), but only one of 12 dense datasets (0.337 versus 0.211).

These results are consistent with the trust-region constraint (Equation 9) limiting corrections to a mismatched neural prediction when dense data support larger changes. We chose this bound to stabilize inference under sparse sampling (Section 2.2). With highly dense sampling, we expect feems to perform at least as well as dreems, especially if the data generating process is very unlike the simulations that were used to train dreems.

### 3.4 Applications to empirical data

We applied dreems to three previously published datasets: North American gray wolves (*Canis lupus*) (Schweizer et al., 2016), darter fishes (*Allohistium* spp.) (MacGuigan et al., 2026), and blacklegged ticks (*Ixodes scapularis*) (Dong et al., 2025). Each of these publications originally used feems for inference. We first reproduced their results by rerunning feems on each dataset and selecting the best-fitting model using leave-one-out cross-validation over a set of ten *λ* values and ten *λ*_*q*_ values geometrically spaced from 10^*−*3^ to 10^2^ and 10^*−*2^ to 10^2^, respectively (Supplementary Figures S4, S5, and S6). We then applied dreems to each dataset using the program defaults. For the wolf and darter datasets, we used the sparse spatial grid. The blacklegged ticks dataset had significantly more samples, so we used a dense spatial grid with 2,860 nodes.

Running times for both methods are shown in Table 1; both were run on CPU. The feems times include cross-validation, whereas the dreems times cover prediction after training. Across these examples, we observed speedups of roughly two to three orders of magnitude compared to feems, with larger gains observed for graphs with more nodes. These gains reflect the amortization of dreems’ initial simulation and training costs across subsequent analyses.

**Table 1:** CPU execution times for empirical analyses. dreems times include prediction and covariance refinement after training; feems times include cross-validation over ten *λ* and ten *λ*_*q*_ values and the final fit. Simulation and training costs are excluded from dreems times. *N* is the number of individuals. The tick dataset is analyzed at two grid resolutions.

| **Dataset** | **SNPs (N)** | **Nodes** | **Runtime (sec)** |  |
|---|---|---|---|---|
|  |  |  | **DREEMS** | **FEEMS** |
| Wolves | 17,729 (111) | 1,207 | 22.82 | 18,019.73 |
| Darters | 3,333 (97) | 587 | 11.60 | 726.84 |
| Blacklegged ticks | 10,277 (517) | 850 | 11.70 | 1,451.63 |
| Blacklegged ticks | 10,277 (517) | 2,860 | 20.12 | 12,616.90 |

The wolf surface recovers many of the geographic features inferred by feems (Figure 6; Supplementary Figure S4), including lower migration near mountain regions in British Columbia, St. Lawrence Island, and Hudson Bay. The estimates differ across the Canadian Prairies and along Baffin Island, where dreems predicts lower migration. Agricultural development in the Prairies (Danysk, 1995) and localized Arctic gene flow are possible explanations, but these regions also have high predictive SDs (Figure 6b); our inferences cannot rule out either mechanism.

**Figure 6:**
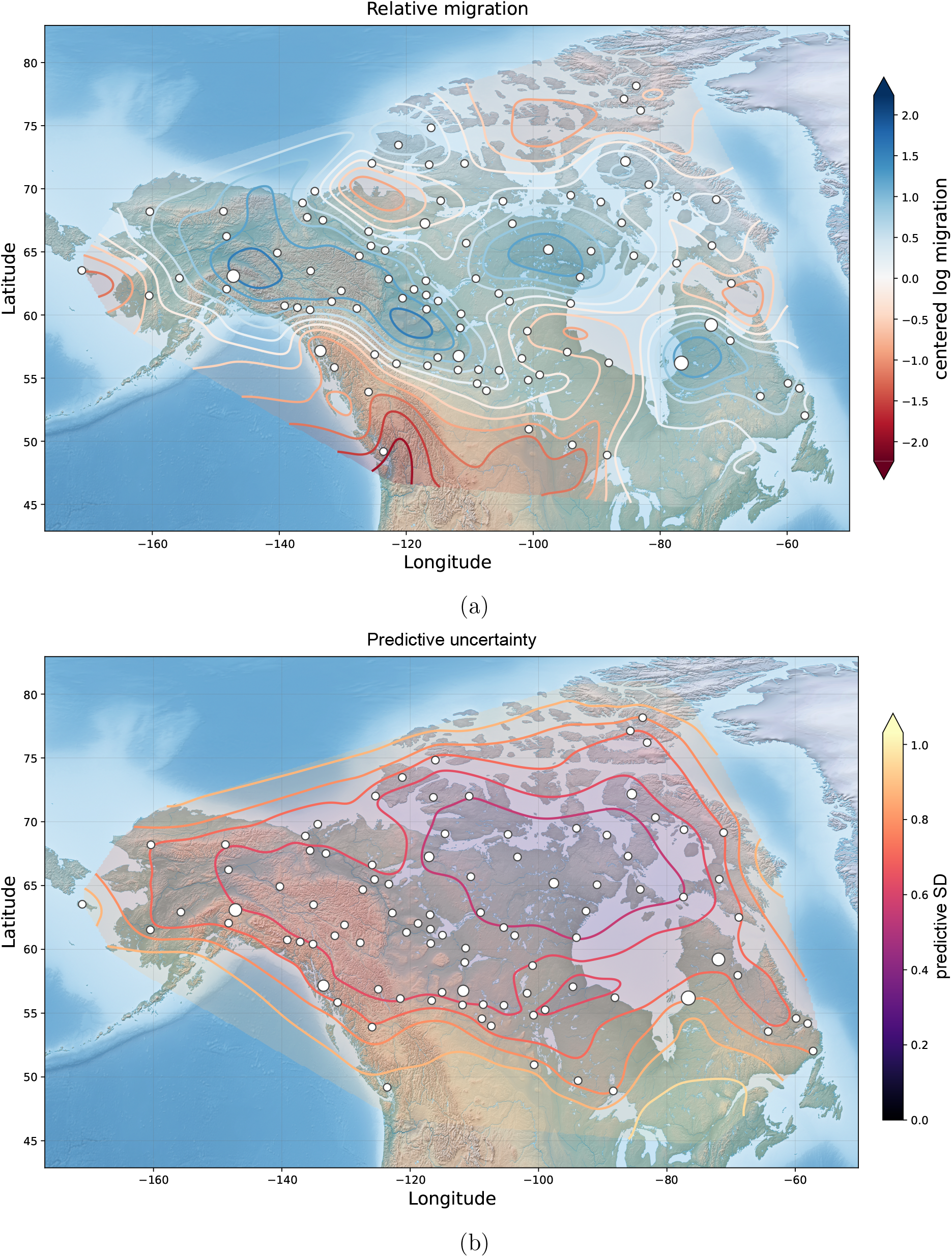
dreems estimates for North American gray wolves on a grid with 1,207 nodes. **(a)** Relative log-migration, with blue indicating higher migration and red lower migration. **(b)** Neural predictive SD, with larger values indicating greater uncertainty. Node sizes indicate sample counts.

The darter surface broadly agrees with feems (Figure 7; Supplementary Figure S5). Both methods infer higher local migration for *A. maydeni* and *A. anas* within their respective river systems and lower migration for *A. cinereum*. dreems also infers reduced connectivity between the Cumberland River *A. maydeni* populations and Duck River *A. anas* populations. feems shows lower migration near some sample clusters, a feature less pronounced in dreems. The shared broad patterns are consistent with the population structure reported by MacGuigan et al. (2026).

**Figure 7:**
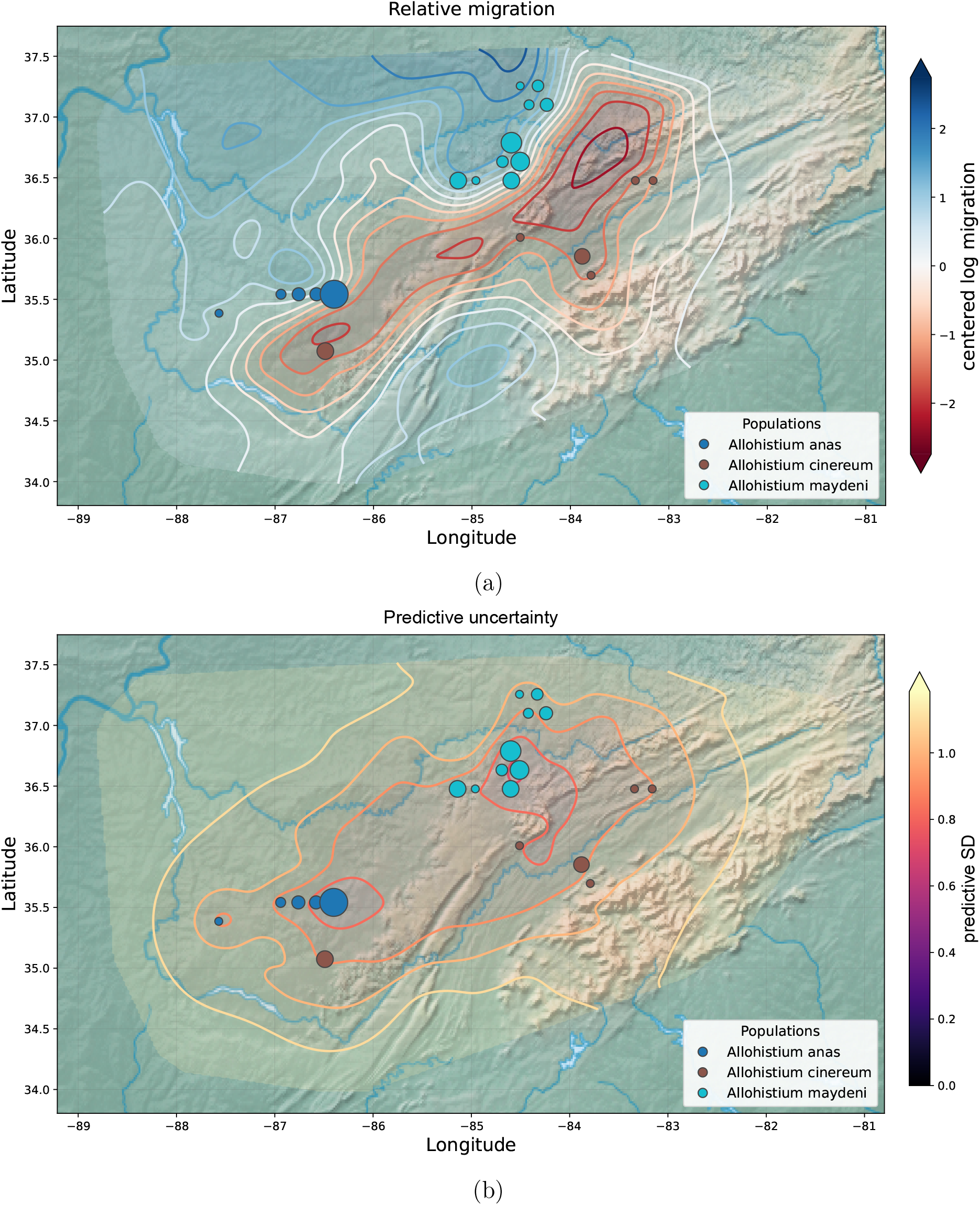
dreems estimates for darter fishes. **(a)** Relative log-migration rate. **(b)** Neural predictive SD.

The blacklegged tick surface shows high connectivity within Michigan’s Upper and Lower Peninsulas (Figure 5). Both methods infer reduced connectivity between Michigan and Wisconsin (Supplementary Figure S6), and dreems also separates the Indiana samples, broadly agreeing with the population structure reported by Dong et al. (2025). High inferred connectivity across Lake Michigan could reflect dispersal mediated by avian hosts, but the surface alone does not distinguish this route from other dispersal histories.

In this example we used a denser grid for dreems than in the published feems analysis, so differences between the displayed surfaces may reflect both grid resolution and inference method. The rapid predictions from dreems make it practical to explore how its estimates change with grid density. On the sparser grid (Supplementary Figure S7), dreems did not completely separate Michigan and Wisconsin. Shared genetic variation among some samples and smoothing across a coarser grid may contribute to this connection. The original study also reported evidence of recent population expansion and cautioned that these populations may not have reached migration–drift equilibrium (Dong et al., 2025), limiting interpretations of the inferred surface.

### 3.5 Sensitivity to grid design

We assessed grid sensitivity by simulating data on an 8 ×12 lattice and analyzing the same observations at four inference-grid resolutions. With dense sampling, the central barrier is recovered on both coarse and fine grids (Supplementary Figure S8). Under sparse sampling, finer grids produce fragmented estimates near sampled locations, while coarser grids recover a more continuous barrier (Figure 8). Even with dense sampling, increasing grid resolution can obscure anisotropic migration (Supplementary Figure S9). These examples emphasize the importance of checking whether inferred features persist across grid resolutions.

**Figure 8:**
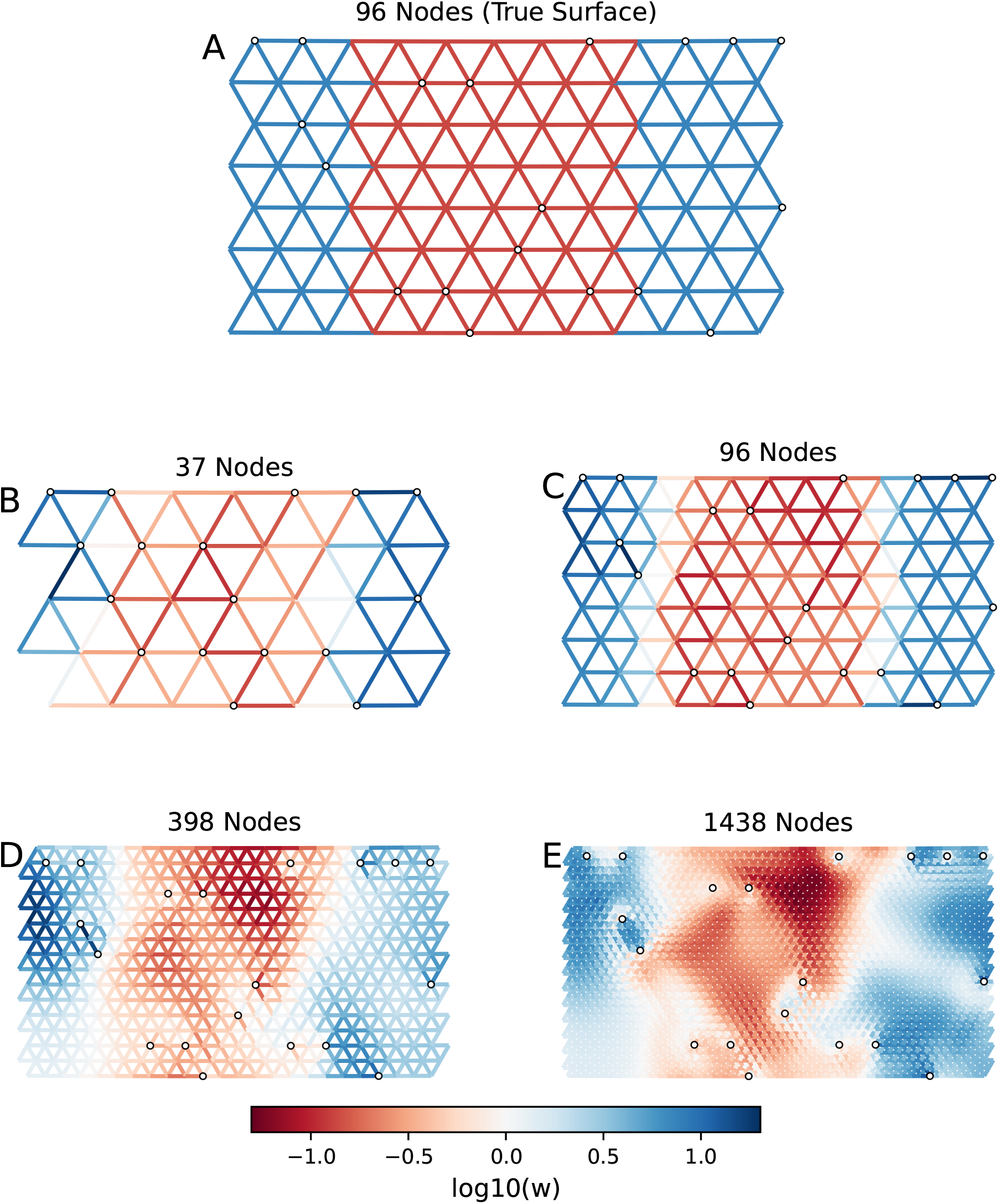
dreems estimates under sparse sampling at four grid resolutions. The true surface, simulated on an 8 ×12 grid with a central migration barrier, is shown in **(a).** Finer inference grids produce fragmented estimates concentrated near sampled locations shown in **(b)-(e)**.

### 3.6 Analyzing polyploids

Finally, we studied whether our method could be used to infer migration surfaces from polyploid genomes, which are common in certain taxonomic groups (e.g. ferns and angiosperms; Hendrickson and Cruzan, 2024; Perrier et al., 2026; Ribicoff et al., 2025). Because of the way our method summarizes its input data, using only deme-wise covariance summaries, we hypothesized that its performance would be relatively agnostic to input ploidy, even though it was trained using exclusively diploid data.

We first confirmed that dreems could accurately recover the main features of migration surfaces from simulated tetraploid and hexaploid genomes (Supplementary Figures S10 and S11). For both ploidies, the central band of reduced migration was recovered under dense sampling and when 80% of nodes were unsampled. With dense sampling, dreems also recovered higher migration along horizontal than diagonal edges in the anisotropic scenario. This directional contrast was weaker under sparse sampling, with greater variation among edges of the same orientation. Note that these analyses used the diploid-trained network without any retraining.

We further evaluated dreems on a real hexploid dataset to examine gene flow in North American yellow birch (*Betula alleghaniensis*) (Cummins et al., 2026). As shown in Figure 9, yellow birch exhibits high gene flow along the Appalachian Mountains and connectivity extending westward into the Great Lakes. These results align with findings reported by Pike and Kern (2022), where yellow birch has high rates of seed and pollen dispersal, facilitating extensive gene flow and connectivity across the migration surface.

**Figure 9:**
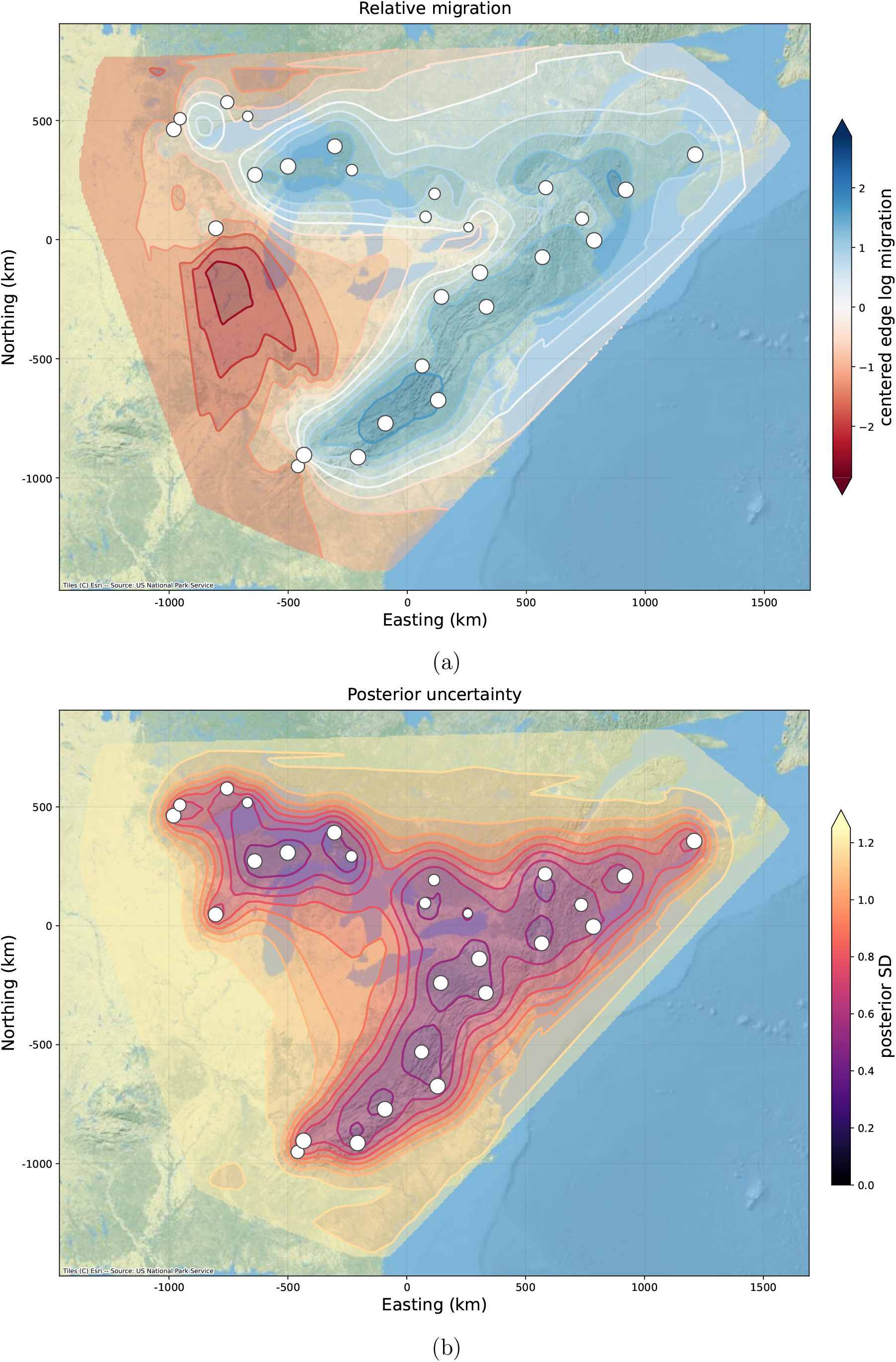
dreems estimates for North American yellow birch. **(a)** Relative log-migration rates. **(b)** Neural predictive SD.

The dreems migration surface reflects the species’ two established glacial lineages: a primary eastern group extending from the Southern Appalachians to New England, and a western group in the Great Lakes region. Although the migration surface shows that these groups are weakly connected, our findings support the presence of two distinct migration clusters. Additionally, hybridization between yellow birch and related sympatric species in the lower and western Great Lakes regions likely differentiates these populations genetically from non-hybridized trees, consistent with reduced migration rates in that region (Thomson et al., 2015). These results also broadly reflect the findings reported by Cummins et al. (2026). The lower migration rates we infer in the Southern Appalachian Mountains and western Great Lakes coincide with regions of lower genetic offset, which quantifies the magnitude of allele frequency shifts required for a population to remain adapted to future climates.

## 4 Discussion

In this paper, we have shown that a neural network trained on simulated data can estimate effective migration surfaces quickly and accurately from genetic covariance. Once trained, the same network can be applied to different datasets, sampling designs, and graphs. This makes it practical to repeat an analysis on several grids, which our empirical results suggest can affect the location and appearance of inferred barriers and corridors. The network also recovered migration patterns from simulated tetraploid and hexaploid data, despite having been trained only on diploid genomes.

We compared dreems with feems because feems is fast and straightforward to apply (Marcus et al., 2021). The largest gains in accuracy occurred when samples or markers were limited, as is often the case in practice. Under these conditions, patterns learned from simulations can help estimate migration in poorly sampled regions.

On the other hand, with sufficiently dense and informative data, we expect likelihood-based methods to perform at least as well as dreems, especially when the data differ sub-stantially from the training simulations. One possibility for improving out-of-distribution performance is to expand the diversity of the training simulations, ensuring that the network encounters a wider range of migration surfaces, sampling designs, and other features. Similarly, it is often possible to fine-tune a general purpose neural network on a smaller set of simulations that more closely match the target dataset (Yosinski et al., 2014). Both strategies could be useful extensions to our method.

Some other additions to the method could further enhance its flexibility and applicability. Adding edges between distant demes would allow us to represent long-range corridors alongside local migration. This requires no change to the neural architecture, which applies to any graph without requiring that edges join geographic neighbors. With suitable training simulations, the network could estimate rates on these additional edges along with rates on the local connections. This would allow us to study long-range genetic similarity, a problem also addressed by feemsmix (Shastry et al., 2025).

Generalizing to asymmetric migration, as in FRAME (Shen and Novembre, 2026), would require more changes. The current network predicts one rate for each undirected edge. To estimate a rate in each direction, it would need to retain the order of the two endpoints and be trained on simulations with asymmetric migration. We would also need to adapt the resistance-based calculations used for initialization and refinement, which assume symmetric migration and can give misleading results under asymmetric gene flow (Lundgren and Ralph, 2019). From a statistical standpoint, inferring asymmetric migration is substantially harder, and how well the genetic summaries used here distinguish migration direction from other forms of variation (e.g. in local population size) remains to be tested.

Another interesting extension is to incorporate environmental and ecological covariates into the model. Habitat, topography, and land use could be included as node or edge features. Simulating migration in relation to these variables would allow the network to learn how they predict connectivity, as explored in landscape-resistance methods such as ResistanceGA (Peterman, 2018). We could also extend the method to estimate migration during different time periods. For example, MAPS uses shared haplotypes to infer recent migration and population size (Al-Asadi et al., 2019). Adding such summaries, or samples collected at different times, could help separate recent changes from longer-term patterns of connectivity.

Finally, neural posterior estimation offers a way to describe uncertainty in whole migration surfaces (Min et al., 2026). Our present estimates give a predictive standard deviation for each edge, but do not describe how errors on different edges are related. Drawing migration and coalescence surfaces jointly would let us examine alternative maps consistent with the data, including how uncertainty in local population size affects the inferred migration.

## Data and Code Availability

All data analyzed in this manuscript were either simulated or previously published (Cummins et al., 2026; Dong et al., 2025; MacGuigan et al., 2026; Schweizer et al., 2016). Code to reproduce our simulated results, as well as an implementation of our method, is available at https://github.com/jthlab/dreems. Documentation and tutorials are available at https://dreems.readthedocs.org.

## Usage of Generative AI

AI was used for code generation, model space exploration, assistance with creation of the figures, and to improve the spelling, grammar, and organization of the manuscript. All generated content was verified by the authors.

## Acknowledgments

This research was supported in part by the National Institute of General Medical Sciences of the NIH under awards number R35GM151145 (JT) and R35GM137919 (GB). The content is solely the responsibility of the authors and does not necessarily represent the official views of the NIH.

