## Supplementary material for "Deep Recovery of Effective Migration Surfaces"

Jiatong Liang[1], Gideon Bradburd[2], Jonathan Terhorst[1]

[1]Department of Statistics, University of Michigan

[2]Department of Ecology and Evolutionary Biology, University of Michigan

### S1 Supplementary methods and figures

#### S1.1 Simulating training data

This section describes how we simulated training and validation data. Graph topology, migration surface, local coalescence rates, and sampling locations varied randomly across training examples, as described in Section S1.1.2. Conditional on these, SNPs were generated using a continuous-time structured coalescent simulator described in Section S1.1.1.

##### S1.1.1 Coalescent simulator

We simulated independent loci using a continuous-time structured coalescent on each spatial graph. Each sampled diploid individual contributes two haploid lineages to its initial deme. A lineage in deme $i$ migrates to a neighboring deme $j$ at rate $m_{ij} = m_{ji}$, and each pair of lineages in deme $i$ coalesces at rate $q_i$. If $k_i$ lineages currently occupy deme $i$, and $k = \sum_i k_i$, the total event rate is

$$\lambda(\boldsymbol{k}) = \sum_i q_i \binom{k_i}{2} + \sum_i k_i \sum_{j:\,(i,j)\in E} m_{ij} + k\theta, \tag{S1}$$

where $E$ is the graph's edge set and $\theta$ is the per-lineage mutation rate. We draw an exponential waiting time with rate $\lambda(\boldsymbol{k})$ and select a migration, mutation, or coalescence event in proportion to its rate. Within the selected deme, a migrating lineage or coalescing pair is chosen uniformly, and a migration destination is chosen in proportion to the outgoing edge rates. Simulation continues until one ancestral lineage remains or until a mutation is encountered.

We use rejection sampling to enforce the minor allele frequency threshold of 0.05. That is, each simulation run is retained only if its derived-allele frequency across all subtended haploids is between 0.05 and 0.95, equivalent to a MAF threshold of 0.05. Note that the MAF can be zero if no mutation occurs before the genealogy reaches its root. Simulation continues until the requested number of usable SNPs is obtained. A fresh genealogy is simulated for each candidate marker, so the loci are unlinked.

The simulator is implemented in Rust, using sparse adjacency lists for migration, restricting event selection to occupied demes, and storing descendant sets as compact bit arrays that are combined at coalescence. For corpus generation, standardized genotype outer products are accumulated directly to obtain $C^{\text{ind}}$ (Equation 1) without needing to store the full genotype matrix.

##### S1.1.2 Motifs

Training graphs were drawn from five motifs covering real and synthetic landscapes. Each motif specifies a graph geometry, migration and coalescence rate fields, and a sampling design.

**Global-landscape simulations** We drew rectangular regions from Natural Earth land maps, varying their location, size, and aspect ratio. We rejected regions centered on water or containing very little land, projected the accepted regions into local planar coordinates in kilometers, and covered them with hexagonal lattices. Water nodes remained in the graph even though samples were taken only on land. Target graph sizes were 220, 450, 900, or 1,400 nodes; grid spacing was adjusted to approach these targets, so the realized counts could differ. Rates combined smooth spatial variation with habitat, water, coastal, and ridge effects (Section S1.2).

Sampling sites were selected uniformly on land, near a random transect, or near a weighted shortest path between two land nodes. We requested samples from 3% to 50% of land nodes, favoring the sparser designs. Each site contained one to eight diploids, with at least two sites per graph and a cap of 384 individuals. This cap could reduce the requested number of sites.

**Simple lattice simulations** These examples used a fixed $8 \times 12$ triangular lattice, choosing among homogeneous, central-barrier, and anisotropic migration with equal probability. The barrier and anisotropic scenarios had fivefold rate contrasts; in the latter, horizontal edges had higher migration. Local coalescence rates varied smoothly across the graph. Each sampled node contained ten diploids. Dense and sparse designs were equally likely: we either sampled every node or sampled each node with probability 0.2.

**Randomized grid simulations** We randomized rectangular grids with heights and widths ranging independently from eight to 36 nodes. We either retained the full grid or removed nodes to form a bay, lake, winding coast, or islands joined by narrow bridges, keeping only the largest connected component. Migration was homogeneous, smoothly varying, or modified by barriers. Local coalescence rates were Gaussian-process draws. Sampling designs either allocated individuals randomly across nodes, gave equal sample sizes to a subset of sites, varied both the number of sites and individuals per site, or sampled every node. Clustered designs concentrated samples around one to eight centers.

**Wildlife-like simulations** These graphs represented corridors and coastlines, with samples concentrated along long paths through the graph. Requested sampling fractions followed the same choices as the global-landscape examples. Migration varied smoothly or included barriers and habitat effects. In habitat-based examples, migration was reduced between dissimilar habitats and along longer edges. Local coalescence rates varied smoothly across nodes.

**Earth-like simulations** These graphs used synthetic landmasses with coasts, islands, bays, or lakes, varying their aspect ratios and choosing target sizes from 80 to 900 nodes. Migration surfaces ranged from homogeneous rates to smooth fields with barriers or habitat effects. Local coalescence rates were Gaussian-process draws, and sampling followed the same designs as the randomized grids.

#### S1.2 Environmental rate fields in global-landscape simulations

For global-landscape graphs, we combined smooth spatial variation with geographic features. Habitat, migration, and local coalescence each started from an independent Gaussian-process field, approximated using random Fourier features. We used a squared-exponential covariance kernel and varied its length scale so that rates could change gradually across a region or over shorter distances. Coordinates were centered and scaled by the region's radius, making these length scales relative to the size of each region.

Habitat quality combined its background field with latitude, proximity to the coast, and land or water status. Coastal influence decayed exponentially with distance from the shoreline, over a distance that varied between simulations. Water lowered habitat scores; latitude and coastal proximity could raise or lower them. The resulting habitat scores were standardized within each graph.

To obtain a migration score for each edge, we averaged the migration background field at its two endpoints and added the habitat and geographic effects. Migration generally increased with the average habitat quality of neighboring demes. We penalized differences between their habitat scores even when both scores were high. We also reduced migration along longer edges, near coasts, and on edges in or crossing water. Water penalties used both endpoints and the edge midpoint, so an edge joining two land nodes could still receive a penalty if its midpoint lay in water.

To add barriers within landmasses, we generated a winding ridge with a random orientation, shape, width, and strength. Edges crossing the ridge received the largest penalty, with weaker penalties on nearby edges.

Local coalescence rates combined their own background field with habitat quality, coastal proximity, and water status. Better habitat generally lowered coalescence rates, corresponding to larger local effective population sizes; water raised them. Coastal effects could act in either direction. We varied the strengths of these effects between simulations, allowing the habitat associations to reverse in some cases. The shared habitat field linked migration and local population size, while the independent background fields allowed each to vary separately.

After combining these effects, we centered and rescaled each log-rate field to its chosen standard deviation. Exponentiating then gave positive migration and coalescence rates with geometric mean one.

#### S1.3 Output weighting and postprocessing

For each edge, we combine two estimates of relative log-migration: the neural decoder's prediction $f_e$ and the centered resistance estimate $u_e$ obtained from Equation 4. The resistance estimate is computed directly from the genetic data, whereas the neural prediction also uses patterns learned from simulations. A learned weight determines how much each estimate contributes to the final blended log-migration estimate $b_e$:

$$b_e = g_e f_e + (1 - g_e) u_e, \qquad g_e \in [0, 1]. \tag{S2}$$

A small neural network computes one scalar $g_e$ for each edge from the decoder's internal features, using a sigmoid output to keep it between zero and one. If the resistance estimate is unavailable for an edge, we use the neural prediction alone.

A second learned weight $h \in [0, 1]$ controls the contrast of the entire surface. We first subtract the mean of the blended estimates, then multiply every edge by this same weight:

$$\mu_e = h \left[ b_e - \frac{1}{m} \sum_{e'=1}^{m} b_{e'} \right], \tag{S3}$$

where $m$ is the number of edges. When $h$ is near one, the centered surface is retained. When $h$ is near zero, log-migration approaches zero on every edge, corresponding to homogeneous relative migration. This *graph gate* can therefore reduce the strength of inferred barriers and corridors without changing their spatial pattern.

Both weights are learned from simulations with known migration rates. The edge weight uses information specific to that edge; the graph weight uses the mean and root mean square of the decoder features across edges, the amplitude of the blended surface, mean predicted log-variance, and the fraction of edges with an available resistance estimate.

#### S1.4 Training objective and optimization

The core training objective combines the edge Gaussian loss (Equation 10) with a local-coalescence Gaussian loss, context-head MSE, and penalties for regression slope, correlation, spatial contrasts, excess error relative to the initializer, and disagreement between subsampled datasets and their more informative counterparts:

$$\begin{aligned} \ell = \ell_{\text{edge}} + 0.5\ell_q + 0.5\ell_{\text{context}} + 0.25\ell_{\text{slope}} + 0.2\ell_{\text{corr}} \\ + 0.35\ell_{\text{contrast}} + 0.5\ell_{\text{init}} + 0.1\ell_{\text{consistency}}. \end{aligned} \tag{S4}$$

- The slope penalty targets a unit regression slope with Huber threshold one.
- The correlation penalty is $1 - r$, where $r$ is the Pearson correlation between the true and predicted edge weights.
- The contrast penalty encourages separation between edges with high and low true migration, targeting at least half the true contrast.
- The initializer penalty is the positive part of the difference between prediction MSE and initializer MSE.
- The consistency loss uses matched simulations with different amounts of data, holding the more informative prediction fixed while updating its subsampled counterpart.

The slope, correlation, and contrast terms are applied only when the true field has root mean square (RMS) greater than 0.05 in training units. Log-variances are clipped to $[-7, 5]$ in those units.

The model fit used 69,632 prepared training examples, in minibatches of eight graphs, Adam with learning rate $5 \times 10^{-5}$, and gradient-norm clipping at one. Validation was performed every 1,000 steps; after 20,000 steps, training stopped if validation loss fails to improve by $10^{-4}$ for 12 consecutive checks. Training stopped at step 38,000 according to these criteria.

#### S1.5 Supplementary figures

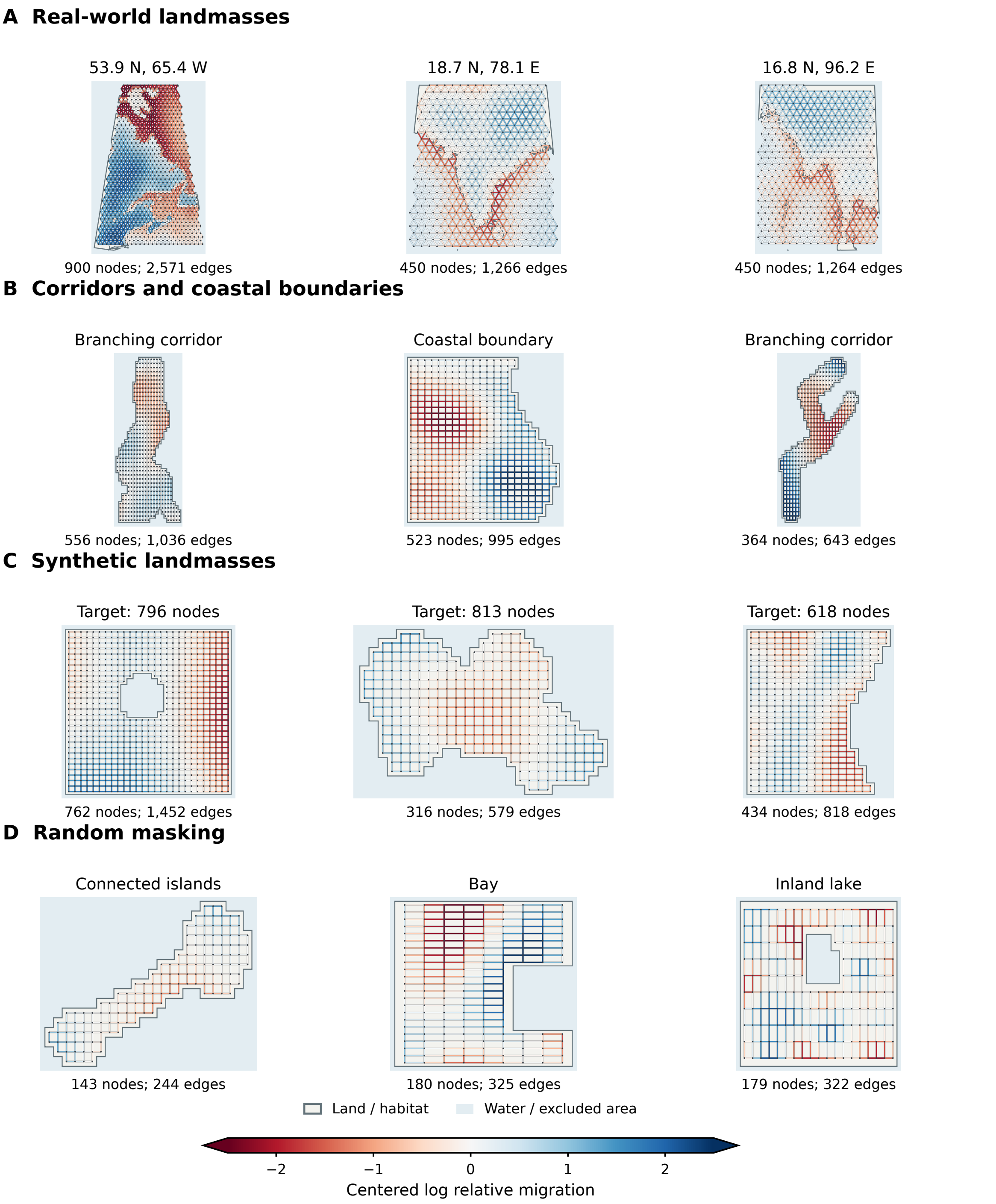


Figure S1: Examples of four randomized graph motifs used to generate training data. **(A)** Real-world landmasses projected onto hexagonal lattices. **(B)** Narrow corridors and coastal boundaries. **(C)** Synthetic landmasses with varied sizes and shapes. **(D)** Randomly masked grids containing connected islands, bays, or inland lakes. Each row shows three examples. Edge colors indicate centered log relative migration rates, with red indicating lower rates and blue indicating higher rates. Pale blue shading indicates water or excluded habitat. Node and edge counts appear below each graph.

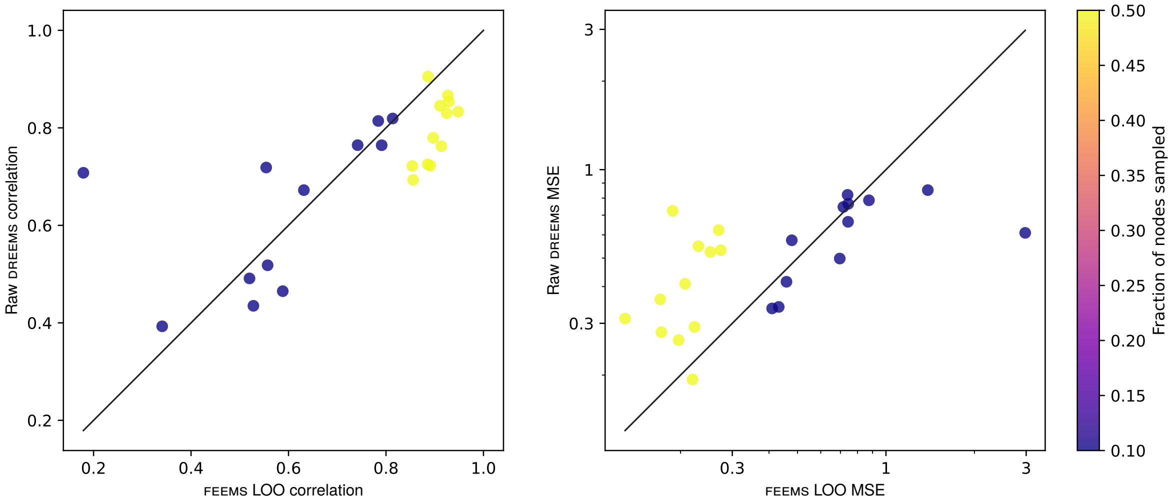


Figure S2: Neural DREEMS predictions before refinement compared with FEEMS using joint leave-one-out cross-validation on the Matérn-1/2 pilot. Each point represents one of 24 datasets from 12 independent migration surfaces, each evaluated under sparse and dense sampling. The left panel shows Pearson correlation with true edge log-migration rates; the right panel shows MSE on logarithmic axes. Points above the identity line in the correlation panel or below it in the MSE panel favor DREEMS. Colors indicate the fraction of nodes sampled: 10% (blue) or 50% (yellow).

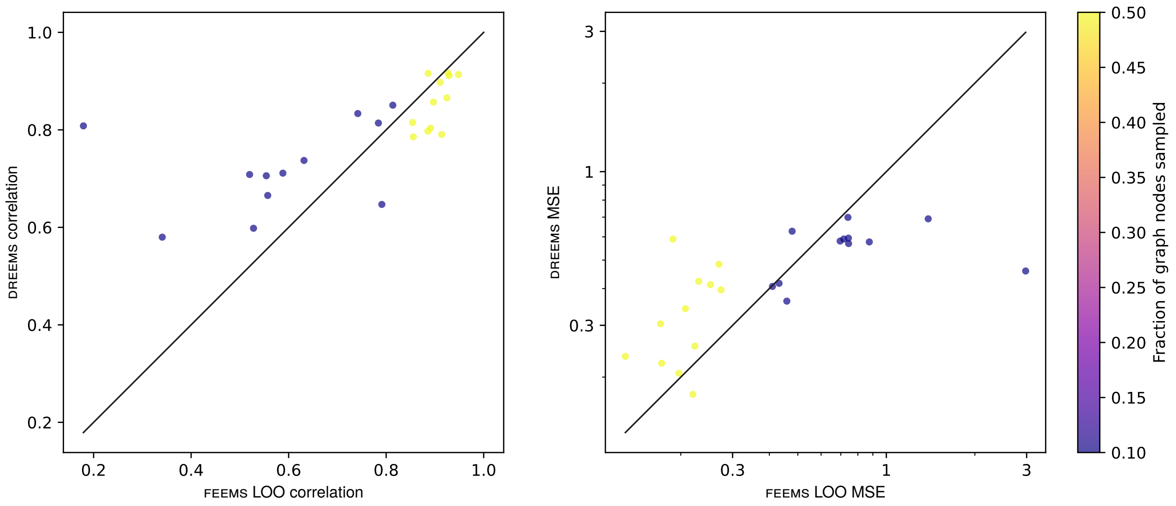


Figure S3: DREEMS with the default covariance refinement compared with FEEMS on the same 24 Matérn-1/2 pilot datasets as Figure S2. Panels, identity lines, and colors follow the same conventions. DREEMS has lower MSE than FEEMS in 11 of 12 sparse datasets, whereas FEEMS has lower MSE in 11 of 12 dense datasets.

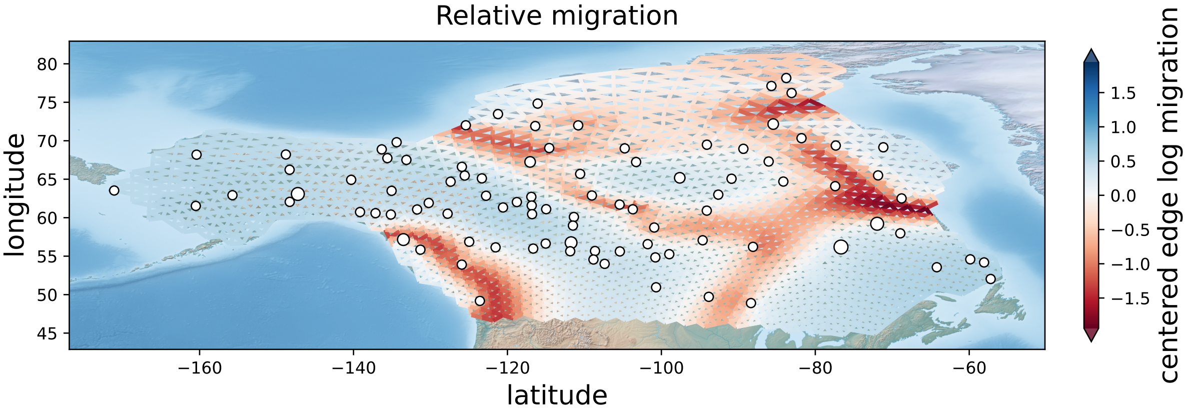


Figure S4: FEEMS estimates of relative migration for North American gray wolves.

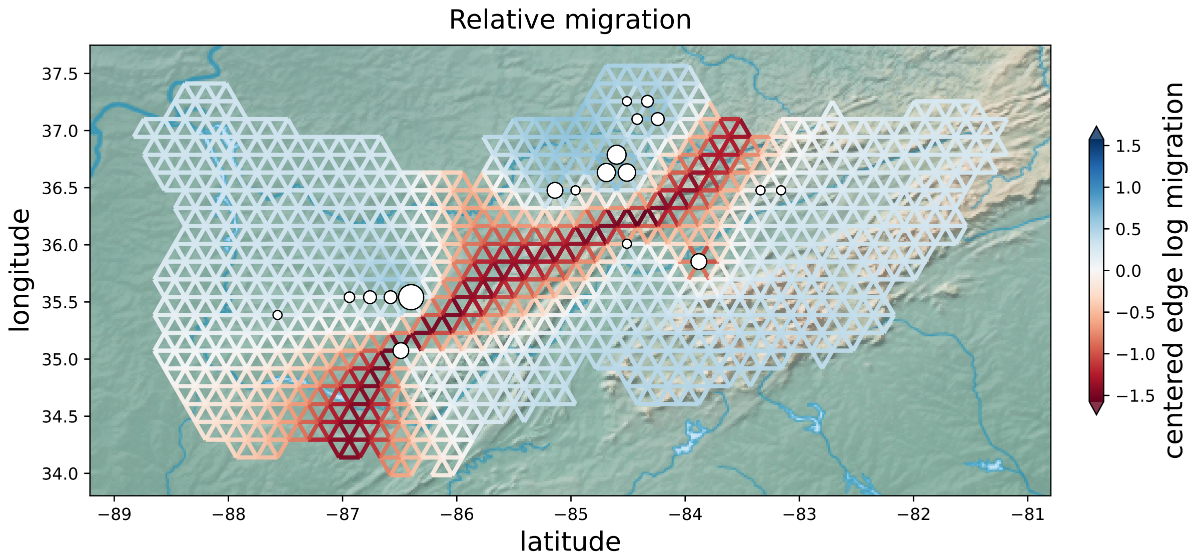


Figure S5: FEEMS estimates of relative migration for *Allohistium* fishes.

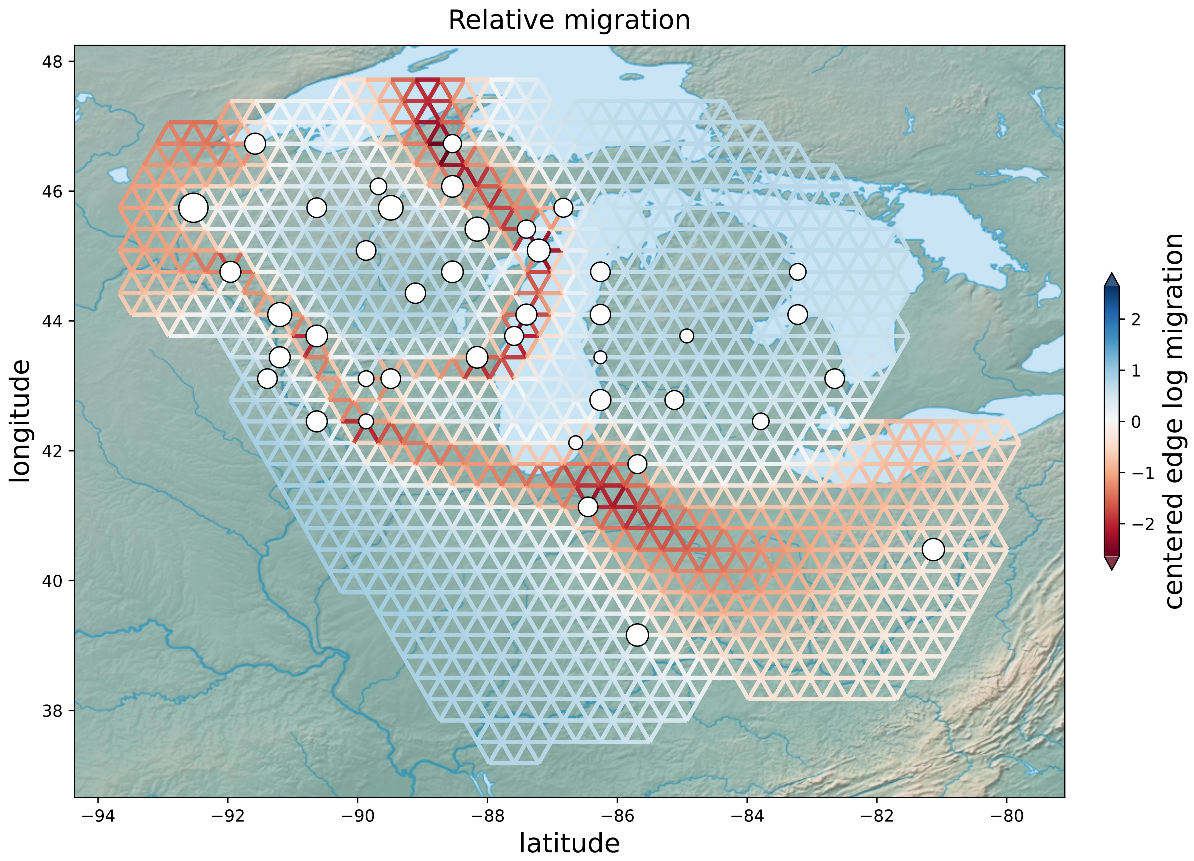


Figure S6: FEEMS estimates of relative migration for blacklegged ticks.

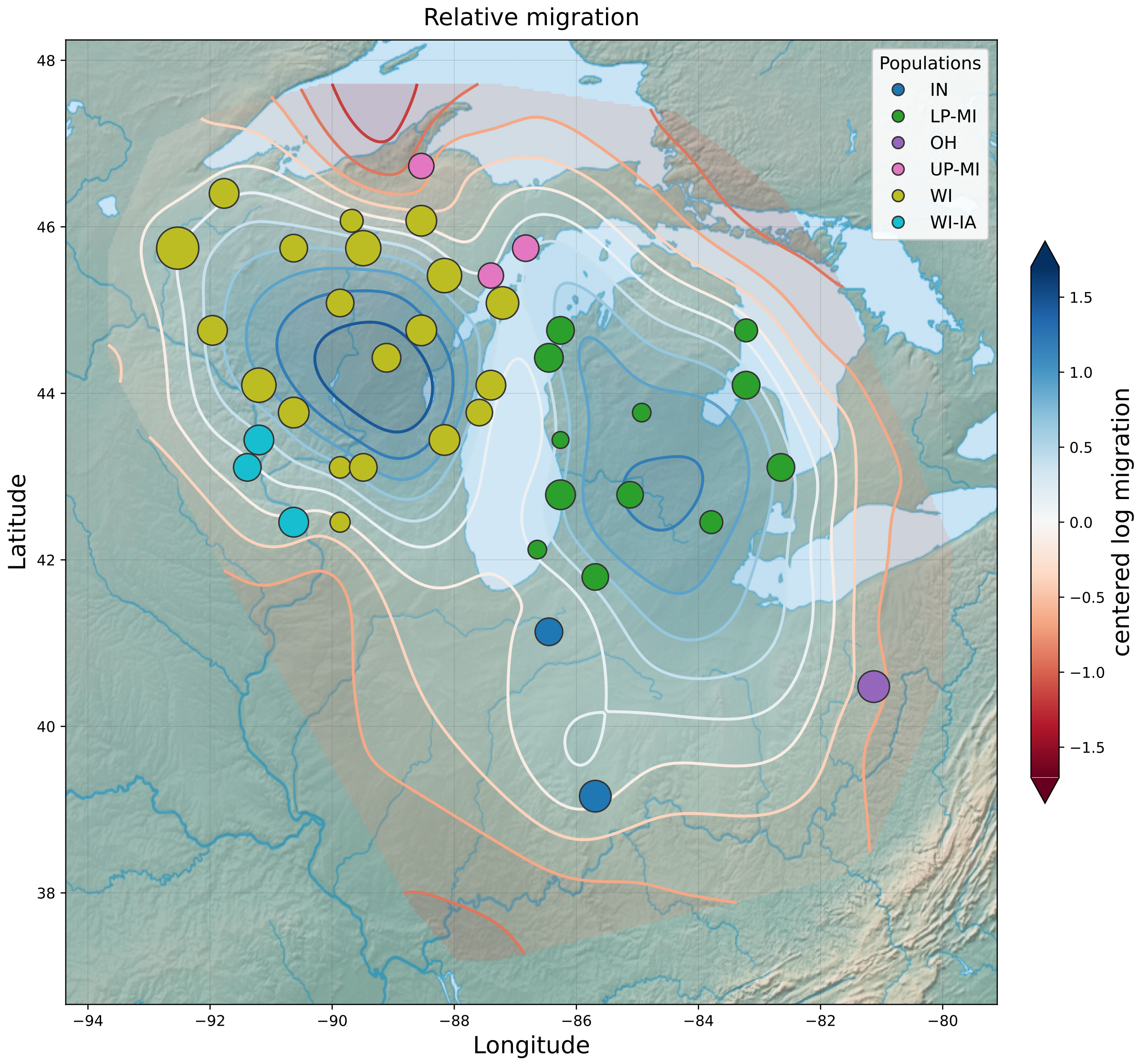


Figure S7: DREEMS estimates for blacklegged ticks on a sparse grid with 850 nodes. The surface retains a connection between Michigan and Wisconsin that is less pronounced on the denser grid (Figure 5).

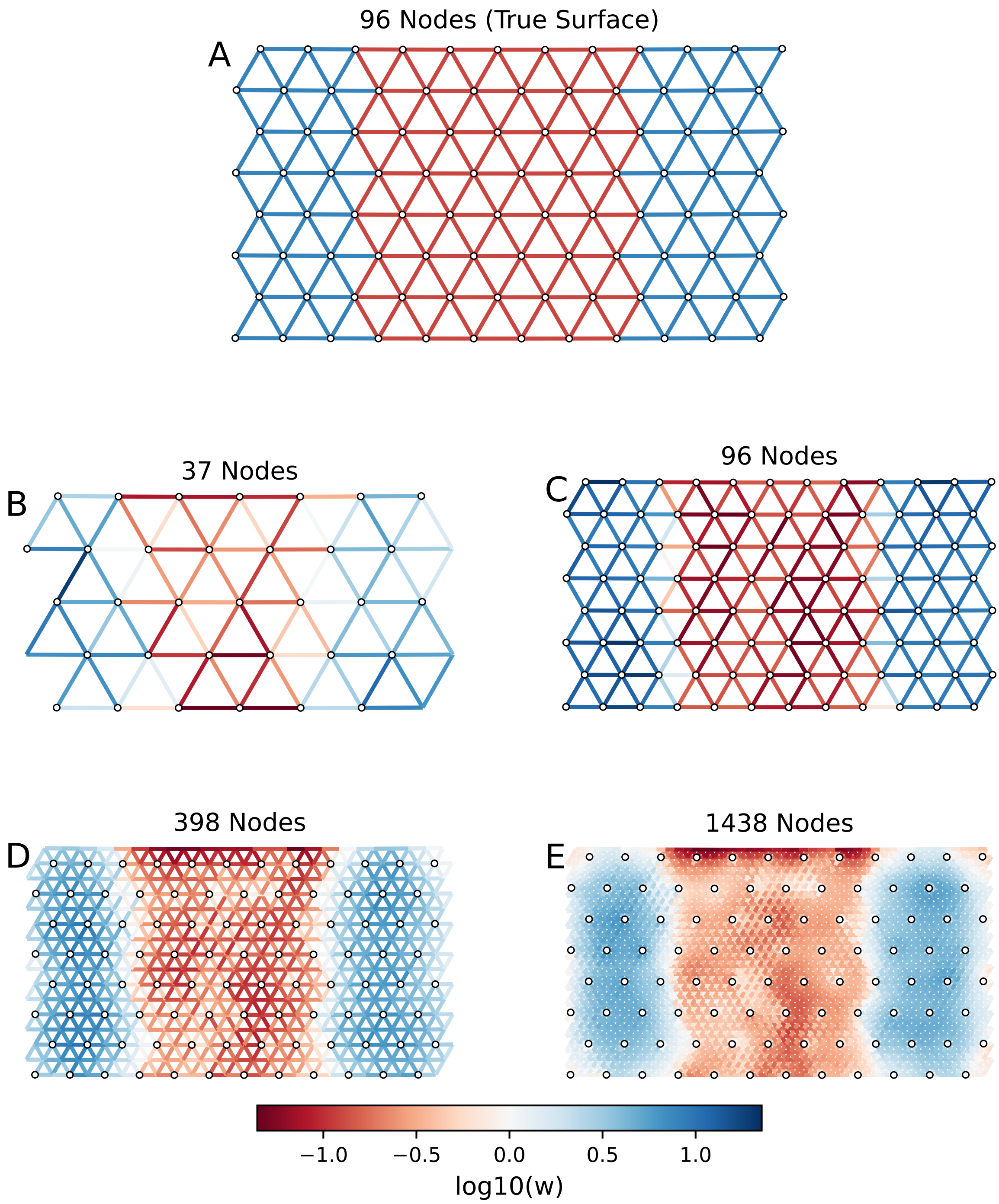


Figure S8: DREEMS estimates under dense sampling at four grid resolutions. The true surface, simulated on an $8 \times 12$ grid, is shown in **(a)**. The central migration barrier is recovered at each tested resolution in **(b)**-**(e)**.

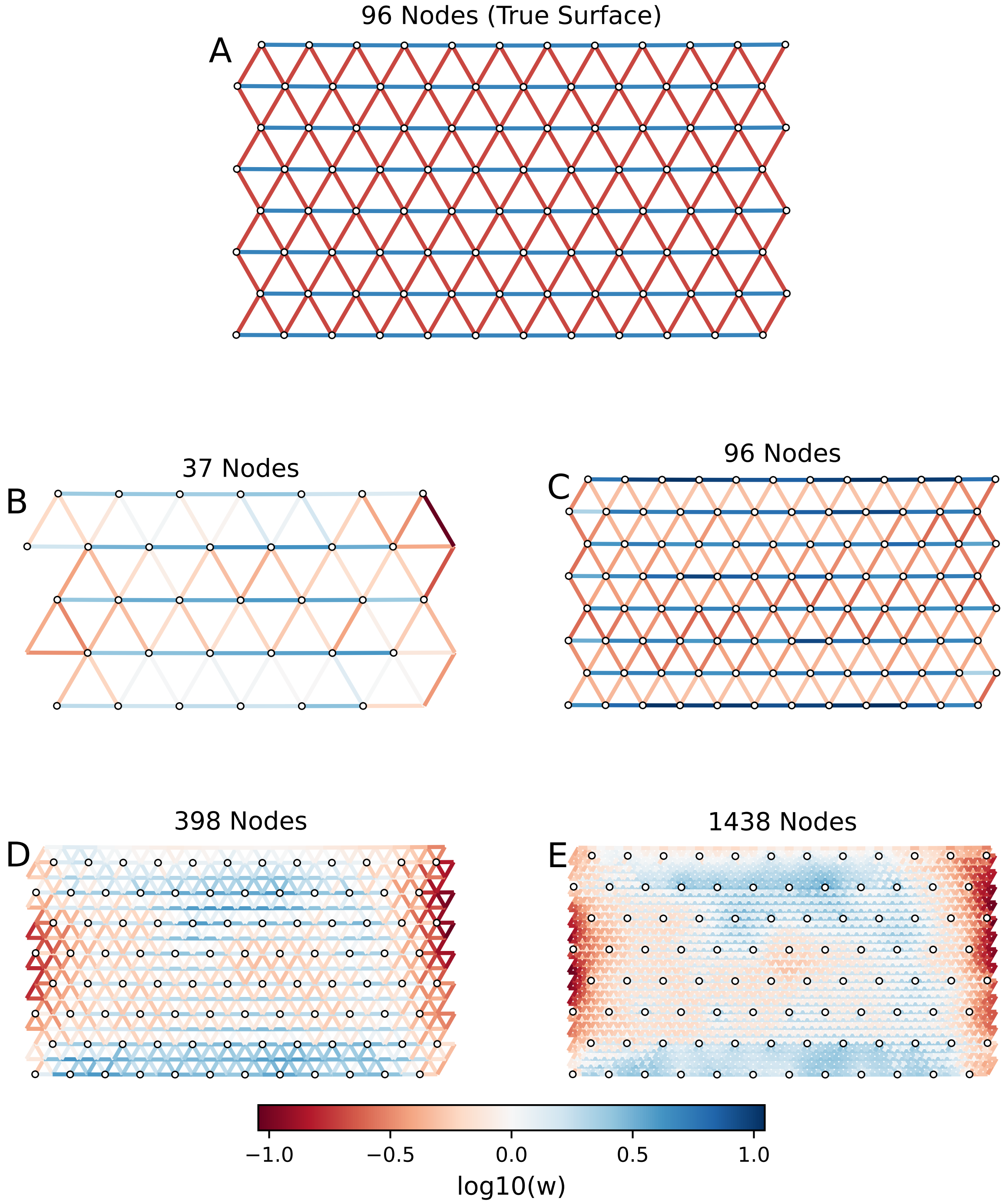


Figure S9: DREEMS estimates of anisotropic migration under dense sampling at four grid resolutions. The true surface, simulated on an $8 \times 12$ grid, is shown in **(a)**. Coarser grids recover higher migration along horizontal edges; finer grids obscure this contrast **(b)**-**(e)**. Migration is symmetric on every edge.

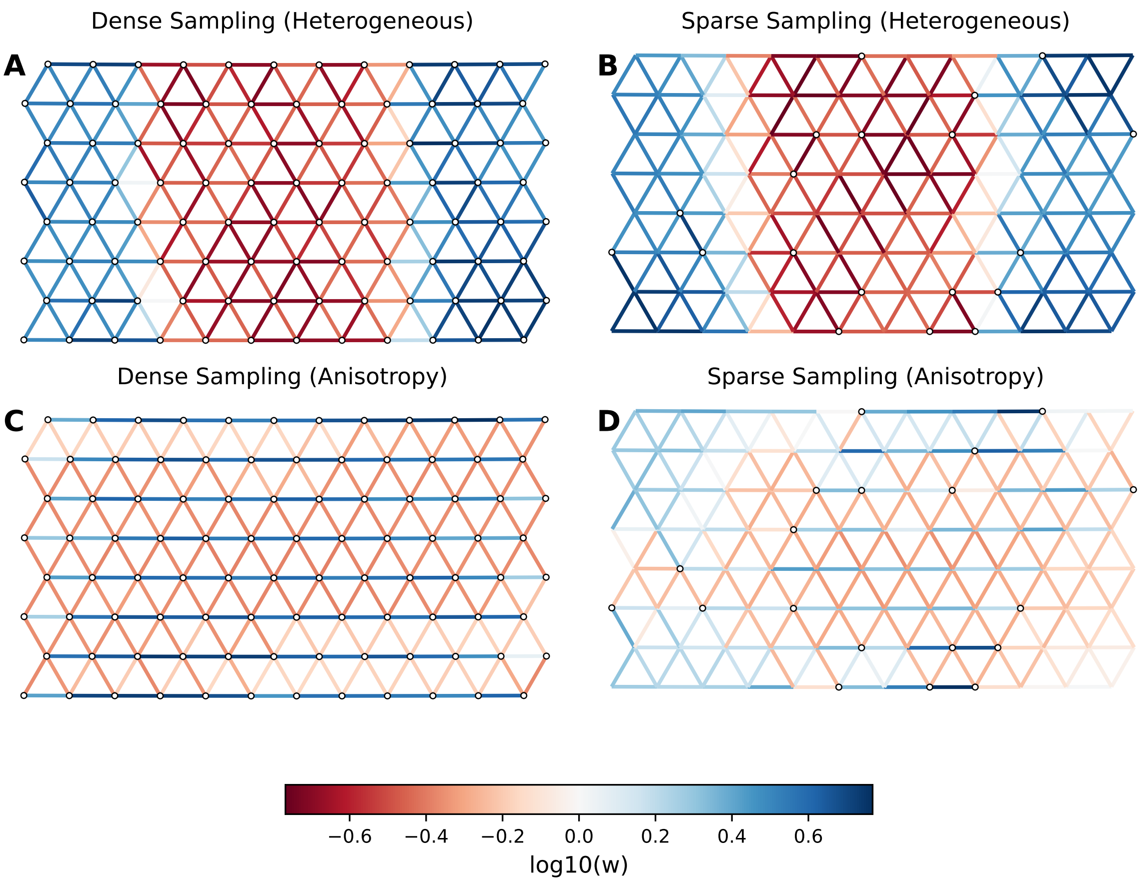


Figure S10: DREEMS estimates from tetraploid simulations. Rows show a central migration barrier **(a)-(b)** and anisotropic migration **(c)-(d)**. Columns show dense sampling (left) and sparse sampling with 80% of nodes omitted (right).

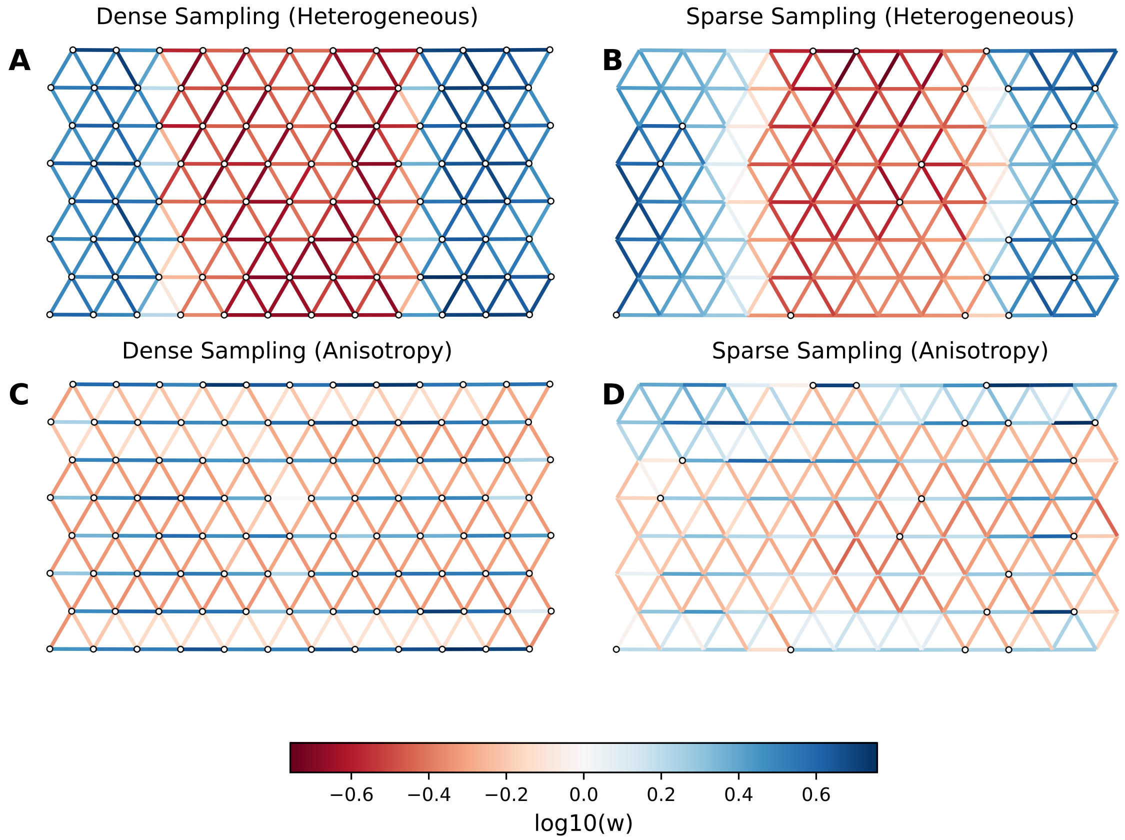


Figure S11: DREEMS estimates from hexaploid simulations. Rows show a central migration barrier **(a)-(b)** and anisotropic migration **(c)-(d)**. Columns show dense sampling (left) and sparse sampling with 80% of nodes omitted (right).
